# Serotonin acts through multiple cellular targets during an olfactory critical period

**DOI:** 10.1101/2024.04.14.589413

**Authors:** Ahana Mallick, Hua Leonhard Tan, Jacob Michael Epstein, Quentin Gaudry, Andrew M. Dacks

**Author notes:** Senior Author: These authors contributed equally (Quentin Gaudry, Andrew M. Dacks).

## Abstract

Serotonin (5-HT) is known to modulate early development during critical periods when experience drives heightened levels of plasticity in neurons. Here, we take advantage of the genetically tractable olfactory system of *Drosophila* to investigate how 5-HT modulates critical period plasticity in the CO2 sensing circuit of fruit flies. Our study reveals that 5HT modulation of multiple neuronal targets is necessary for experience-dependent structural changes in an odor processing circuit. The olfactory CPP is known to involve local inhibitory networks and consistent with this we found that knocking down 5-HT7 receptors in a subset of GABAergic local interneurons was sufficient to block CPP, as was knocking down GABA receptors expressed by olfactory sensory neurons (OSNs). Additionally, direct modulation of OSNs via 5-HT2B expression in the cognate OSNs sensing CO2 is also essential for CPP. Furthermore, 5-HT1B expression by serotonergic neurons in the olfactory system is also required during the critical period. Our study reveals that 5-HT modulation of multiple neuronal targets is necessary for experience-dependent structural changes in an odor processing circuit.

## INTRODUCTION

In early postnatal life of animals, all sensory systems exhibit heightened levels of plasticity and circuit refinement in response to environmental stimuli in a specific time window called critical periods. Critical period plasticity (CPP) provides an excellent readout to assess how sensory experiences shape circuits early in life at the cellular and molecular level.^1–3^ Along with sensory experiences, neuromodulators like 5-HT also play an important role in shaping CPP. Initial experiments indicating the role of 5-HT in modulating sensory critical periods was observed in both the visual and somatosensory cortices.^4–7^ Similarly, 5-HT has also been shown to modulate the early postnatal development of limbic circuits such as the pre-frontal cortex and disruption in serotonergic signaling in this circuit have been linked to an increased risk for behavioral and cognitive deficits in adults.^8–11^ Thus, 5-HT targets multiple effector regions during early development to facilitate proper brain development and function in adults. However, 5-HT can activate several receptor subtypes expressed by distinct cell types within a network, so the cellular mechanisms by which 5-HT impacts CPP can be difficult to identify.

We took advantage of the wealth of transgenic tools and foundational work in the olfactory system of *Drosophila* to determine how 5-HT can impact different circuit mechanisms within an olfactory critical period. The organization of the fruit fly olfactory network is similar to that in mammals^1,12^ in that olfactory processing begins upon odor binding chemoreceptive proteins localized at the dendrites of olfactory sensory neurons (OSNs).^13–16^ All OSNs expressing the same complement of chemoreceptive proteins project to a distinct glomerulus to form an olfactory map within the primary olfactory center, the antennal lobe (AL).^17^ OSNs synapse upon second-order projection neurons that project onto higher order olfactory centers in the mushroom body (MB) and lateral horn (LH). The cellular and molecular mechanisms of structural plasticity observed in olfactory CPP in the fruit fly is well known.^18–27^ Most CPP studies in *Drosophila* were conducted using either CO_2_ or ethyl butyrate exposure as sensory stimuli, or both. With either of the odors, chronic exposure during the critical period induced structural plasticity in the relevant glomerulus causing a change of either an increase or a decrease in the glomerular volume. At the circuit level, the structural plasticity resulting in an increase in glomerular volume is manifested through an increase in the number of LN and PN arbors innervating the glomerulus while the total number of neurons remains intact.^19–21,27^ Glomerular volume decrease, on the other hand is caused by retraction of the OSN axon fibers upon chronic ethyl butyrate exposure in a specific glomerulus.^23^ These studies demonstrated that the olfactory CPP in Drosophila shares many of the same molecular mechanisms as visual CPPs.^2,3,20–23,27–30^ In both cases, CPP involves GABAergic and glutamatergic signaling, Ca2+/Calmodulin dependent adenylate cyclase and cAMP response element binding protein (CREB) dependent gene transcription.^20,21,31^ However, while 5-HT neuromodulation has been shown to be required for visual critical periods in mammals^5–7,32–35^, it has not been studied with respect to CPP in any olfactory system.

In this study, we investigate how 5-HT modulates olfactory critical periods and we focused on the behaviorally relevant^36,37^ CO_2_ sensing circuit in *Drosophila*. Since the V glomerulus is exclusively dedicated to respond to CO_2_ and the CPP mechanisms is already known for this glomerulus, it is ideal for studying the effect of serotonergic modulation. We performed cell-type specific genetic manipulations of the serotonergic system to identify where 5-HT is required during odor-evoked structural plasticity in the olfactory circuit. Our results show that during the critical period, 5-HT modulates distinct cell types in the AL via activation of different 5-HT receptor subtypes. We thus identified cell types where serotonergic modulation may be interacting with the previously described mechanisms of CPP to modulate structural plasticity during CPs.

### Materials and Methods

#### Fly rearing and maintenance

All *Drosophila* lines were raised in sparse cultures on cornmeal, yeast, dextrose medium ^38^ at 25°C in a 12hour light/dark cycle unless otherwise noted. The fly lines used in this study can be found in Table 3 and supplementary Table S1. We used female flies for our studies, except where specifically mentioned. Flies used in the CO_2_ and air exposure experiments were raised at 25°C until the 4-day old pupae stage and at 23°C until they were sacrificed.

#### Odor exposure

We employed previously established odor exposure protocol to induce critical period plasticity in the flies^21,23^ Briefly, 4-day old pupae of relevant genotype were collected into separate vials based on odor-exposure. A fine mesh cheesecloth was secured at the opening of the fly vial to ensure free gaseous exchange. Fly vials were placed in either a temperature-controlled CO_2_ or regular incubator at 23°C on 12-hour light/dark cycles. Eclosed flies were transferred into clean vials 18-21 hours after the start of odor exposure until day 5 post eclosion when they were collected for dissection and immunohistochemistry. For each of the odor exposure experiments, pupae for all the genotypes including control and Canton-S (wild-type) were collected on the same day and odor exposure started at the same time. Pupae of the same genotype were collected from the same stock bottle and separated into odor exposed (CO_2_) and control (air) vials to ensure consistency in food composition, availability, and conditions. During odor exposure both male and female flies of a specific genotype were present in each vial. Only female flies were collected after cold anesthetizing the flies at the end of the experiment right before dissection.

### Plasmid preparation for CRISPR/Cas9 knock-in

The GFP_11_-HA was synthesized de novo by Twist Bioscience, CA and the coding sequence was knocked-in the fly genome using CRISPR/Cas9 via homology-directed repair (HDR). Two plasmid DNAs were prepared for each knock-in preparation: the guide RNA (gRNA) plasmid and the donor plasmid. The gRNA plasmid was prepared following a previously established protocol^39^. Briefly, a pair of 24 nucleotide oligos were commercially synthesized as for normal unsalted primers (Eurofins Genomics). For each oligo, the first 4 nucleotides (TTCG for the sense strand and AAAC for the antisense strand) at the 5’ end formed the overhanging sequence after reannealing. These overhanging nucleotides were compatible with the Bbs I digestion sites of the plasmid pU6b, while the remaining 20 nucleotides carried the sense or antisense strand of the target sequence for the insertion site in the genome. The 20-nt target sequences (sense) were identified close to the C-terminus of the 5-HTR coding region using an online tool FlyCRISPR ^40^ for 5-HT receptor (5-HTR) genes, as listed in **Table 1**. The backbone for gRNA expression, pU6b, was digested with Bbs I and ligated with the reannealed 24-nt oligo pairs with T4 DNA ligase. To prepare the donor plasmid, approximately one kilo basepair of DNA upstream of the insertion site was PCRed as the left arm for homology directed repair (HDR). Similarly, about one kilo basepair of 5-HTR DNA in the downstream of the insertion site was PCRed as the right arm for HDR. The encoding sequence of GFP_11_-HA was then inserted in between both arms via overlap extension PCRs. The resultant PCR product was then inserted into pTwist Amp plasmid (High copy, Twist Bioscience HQ, CA) via restriction digestion and ligation with Rapid DNA Ligation Kit (Thermo Fisher Scientific, Cat. # K1422). For both plasmids, the ligation mixture was then used for transformation of *E. coli* strain DH5α (Thermo Fisher Scientific, Cat. # EC0112). Single colonies of DH5α were used for inoculation of 10 mL mini cultures with LB medium supplemented with 100 μg/mL ampicillin (Thermos Fisher Scientific, Cat. # J60977.06). After overnight incubation with vigorous shaking, plasmid DNA was prepared from the culture using the NucleoSpin Plasmid Mini kit (MACHEREY-NAGEL, Cat. # 740588.250). The purified DNA was digested with restriction enzymes for quality control and subject to sequencing analysis for verification.

**Table 1.**
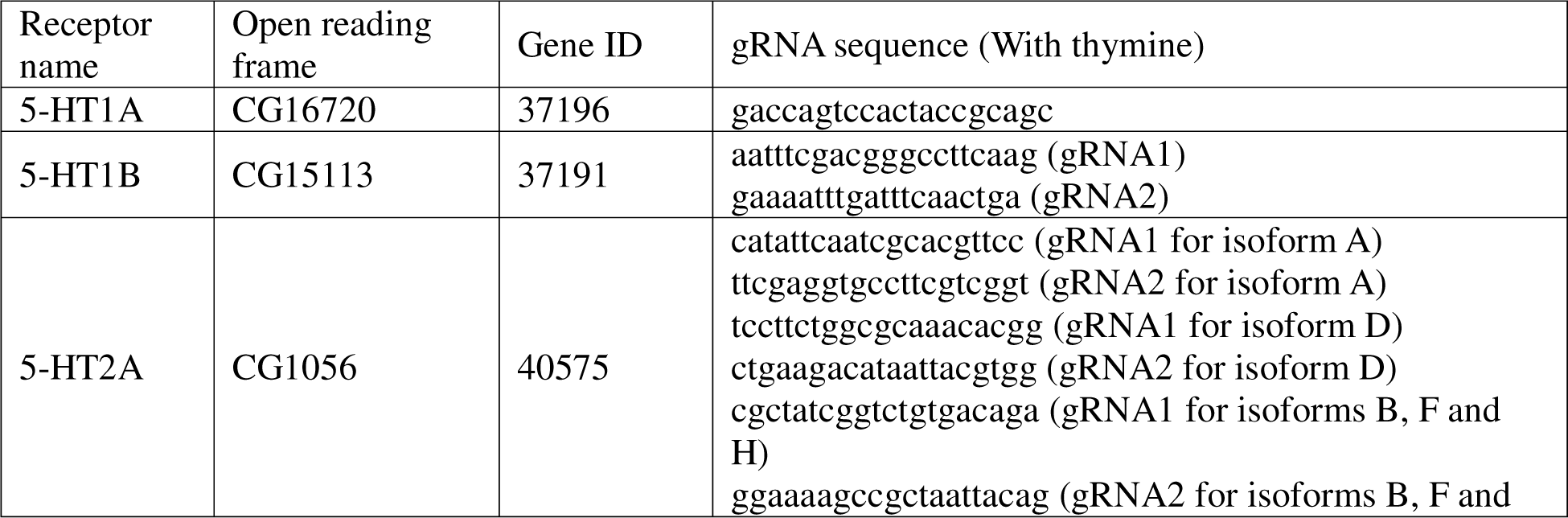
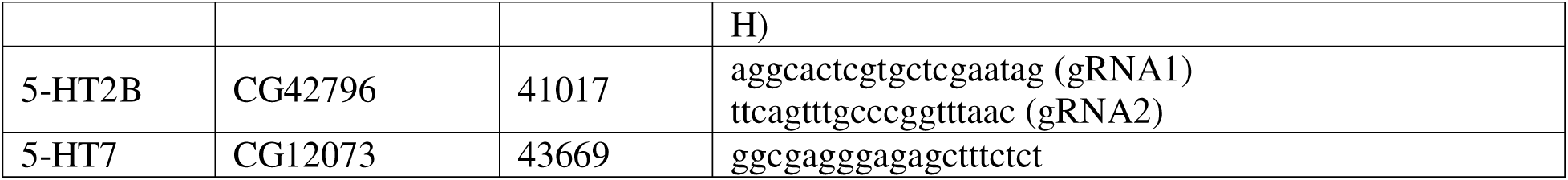
Coding sequences of guide RNA (gRNA) spacers for CRISPR/Cas9 knock-in of GFP11-HA in 5-HTR genes.

### Extraction of genomic DNA

To extract the genomic DNA from adult flies or larvae for generating PCR templates, 2 flies were smashed in 50 μL of Squishing buffer (10 mM TrisCl with pH 8.2, 1 mM EDTA, 25 mM NaCl and 200 μg/mL freshly added Proteinase K) with a 200-μL pipette tip in a PCR tube. The mixture was incubated at 50 degrees Celsius for 30 minutes before inactivated at 95 degrees Celsius for 5 minutes. After it cooled down, 0.5 to 1 uL of the supernatant was used for the PCR template.

### Generation of fly lines

To generate the 5-HTR-GFP_11_-HA transgenic lines, the gRNA plasmid and the donor plasmid was co-injected with a 1:1 mass ratio into fly embryos by Rainbow Transgenic Flies, Inc. (California, USA). The genotype of the embryos was *y,sc,v; {nos-Cas9}attP2* for 5-HT1A and -1B, or *y,sc,v; {nos-Cas9}attP40/CyO* for 5-HT2A, -2B and -7. Depending on which chromosome the 5-HTR gene is on, the candidate flies were individually balanced with Cyo (for 5-HT1A and -1B) by crossing with *w1118; Cyo/Sco*, or *Tm6b, Tb, Hu* (for 5-HT2A, -2B and -7) by crossing with *w1118; Tm6b, Tb, Hu/MKRS*. The nos-Cas9 transgene was finally removed by selecting progeny against red eyes.

Screening for successful knock-in was performed using PCR with a pair of primers targeting respectively the upstream and the downstream of the inserted sequence in the genomic DNA extracted from the candidate transformants. For PCR template, the genomic DNA was extracted from the daughter flies of individual injected embryos. Immunostaining against the HA-tag was used as the second step of verification.

### Immunostaining and confocal imaging

All immunohistochemistry was performed using a standard protocol as previously described unless otherwise noted ^38,41^ For volume measurement experiments, the brains were incubated in the mounting medium for 1 hour before imaging to allow equilibration^42^. For imaging HA and split-GFP, the immunohistochemistry protocol was slightly modified. Briefly, the flies were anesthetized in a glass vial on ice for 1 min. The brains were dissected in PBS and fixed in 4% formaldehyde (37%, diluted in PBS) at room temperature. After three times of washing each for 15 min with 0.2% PBST, i.e., PBS supplemented with 0.2% (v/v) Triton X-100 (Thermo Fisher Scientific, Cat. # A16046.AE), the brains were blocked in 10% normal goat serum (NGS; Thermo Fisher Scientific, Cat. # PCN5000) at room temperature for 1 - 2 hours. Next, the brains were incubated with primary antibodies diluted in 0.2% PBST supplemented with 5% NGS at 4° for 3 to 4 days. Then the brains were washed with 0.2% PBST three times each for 15 min, and subject to incubation with secondary antibodies diluted in 0.2% PBST supplemented with 5% NGS in a dark environment at 4° for 1 day. Table 2 lists all the antibodies used in this study. After washing with 0.2% PBST four times each for 10 min, the brains were mounted on glass slides (VWR, Cat. # 16004-422) with VECTASHIELD Antifade Mounting Medium (Vector Laboratories, Cat. # H-1000-10) and covered with glass coverslips (VWR, Cat. # 48366-089). To prevent the brains from being smashed, two smaller coverslips (VWR, Cat. # 48366-045) were placed as spacers between the glass slide and the covering coverslip with one on each side. The coverslips were fixed with a few drops of nail polish on the edge. All samples were scanned with a confocal microscope in the institution imaging facility: ZEISS LSM980, ZEISS LSM 710 or PerkinElmer Spinning disk. Brains used in the same experiment were imaged on the same day, under the same microscope and image acquisition settings.

**Table 2.**
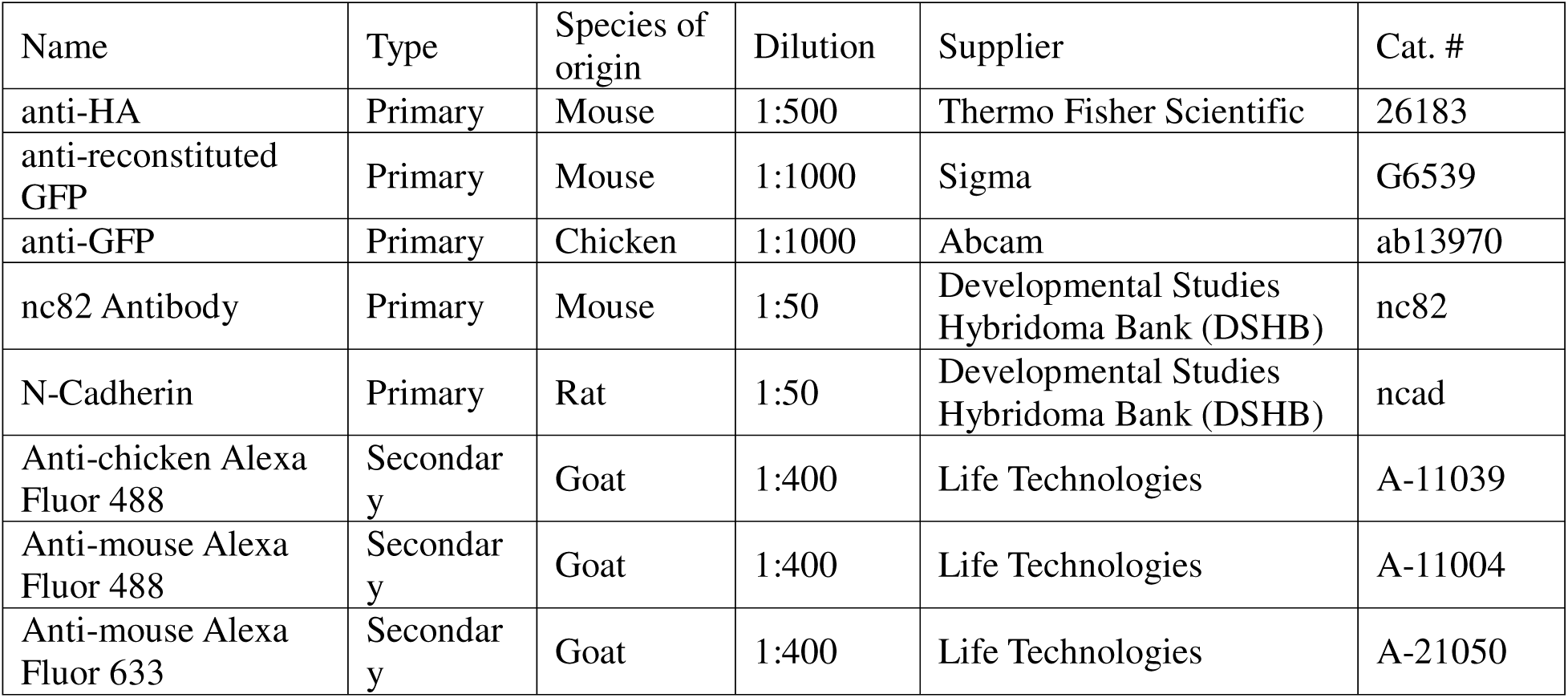
List of Antibodies used in this study.

| Name | Type | Species of origin | Dilution | Supplier | Cat. # |
| --- | --- | --- | --- | --- | --- |
| anti-HA | Primary | Mouse | 1:500 | Thermo Fisher Scientific | 26183 |
| anti-reconstituted GFP | Primary | Mouse | 1:1000 | Sigma | G6539 |
| anti-GFP | Primary | Chicken | 1:1000 | Abcam | ab13970 |
| nc82 Antibody | Primary | Mouse | 1:50 | Developmental Studies Hybridoma Bank (DSHB) | nc82 |
| N-Cadherin | Primary | Rat | 1:50 | Developmental Studies Hybridoma Bank (DSHB) | ncad |
| Anti-chicken Alexa Fluor 488 | Secondary | Goat | 1:400 | Life Technologies | A-11039 |
| Anti-mouse Alexa Fluor 488 | Secondary | Goat | 1:400 | Life Technologies | A-11004 |
| Anti-mouse Alexa Fluor 633 | Secondary | Goat | 1:400 | Life Technologies | A-21050 |

**Table 3.**
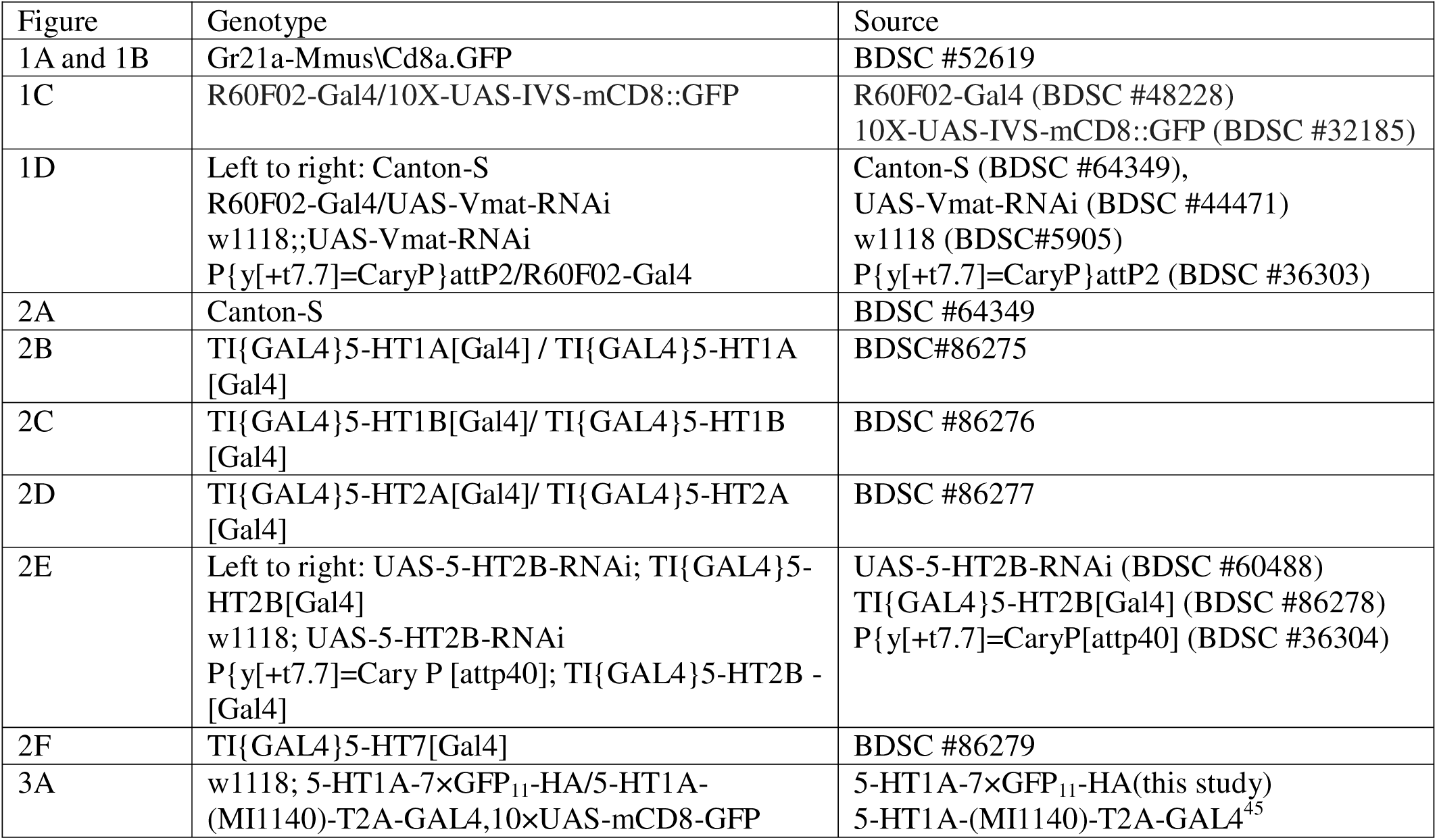
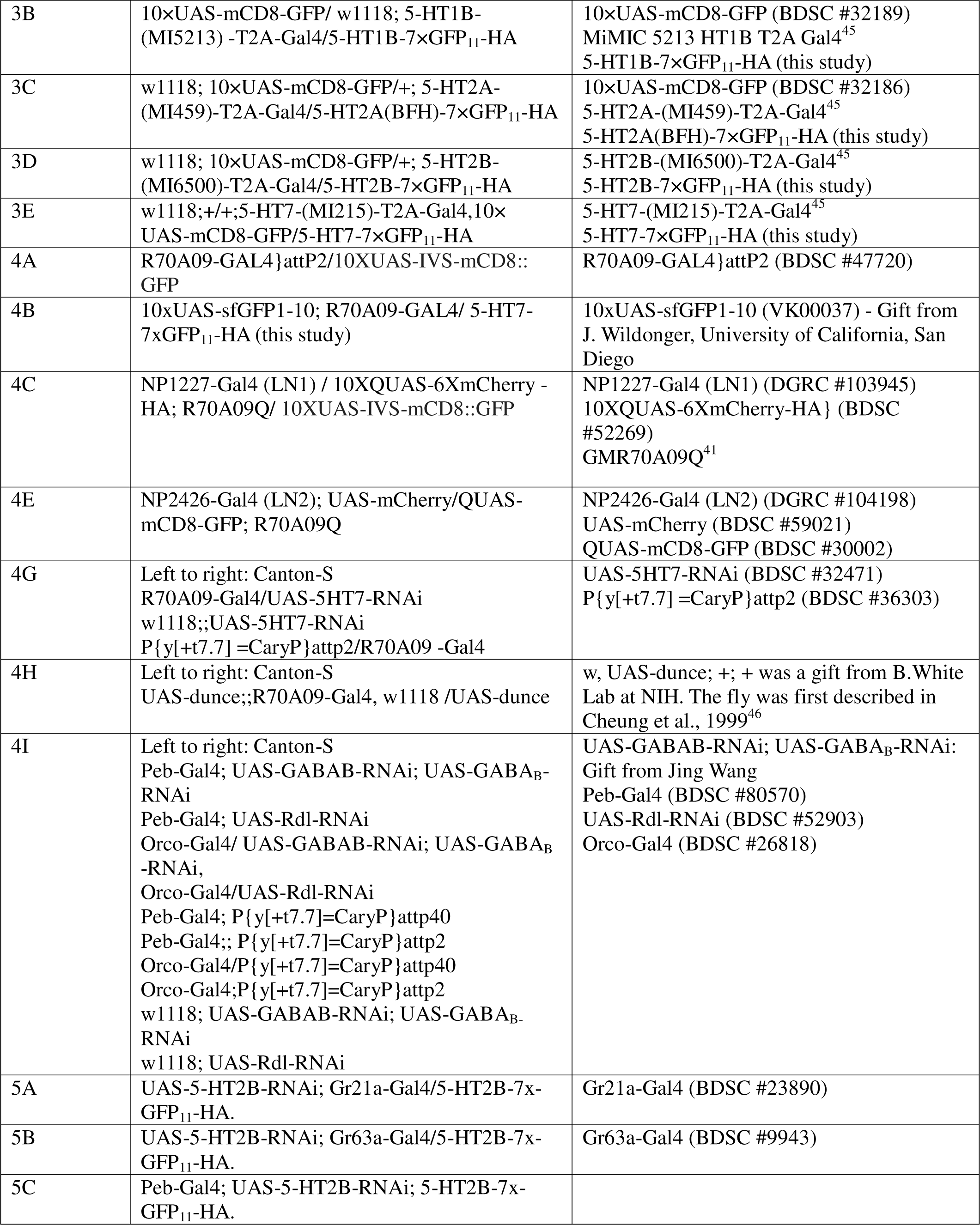
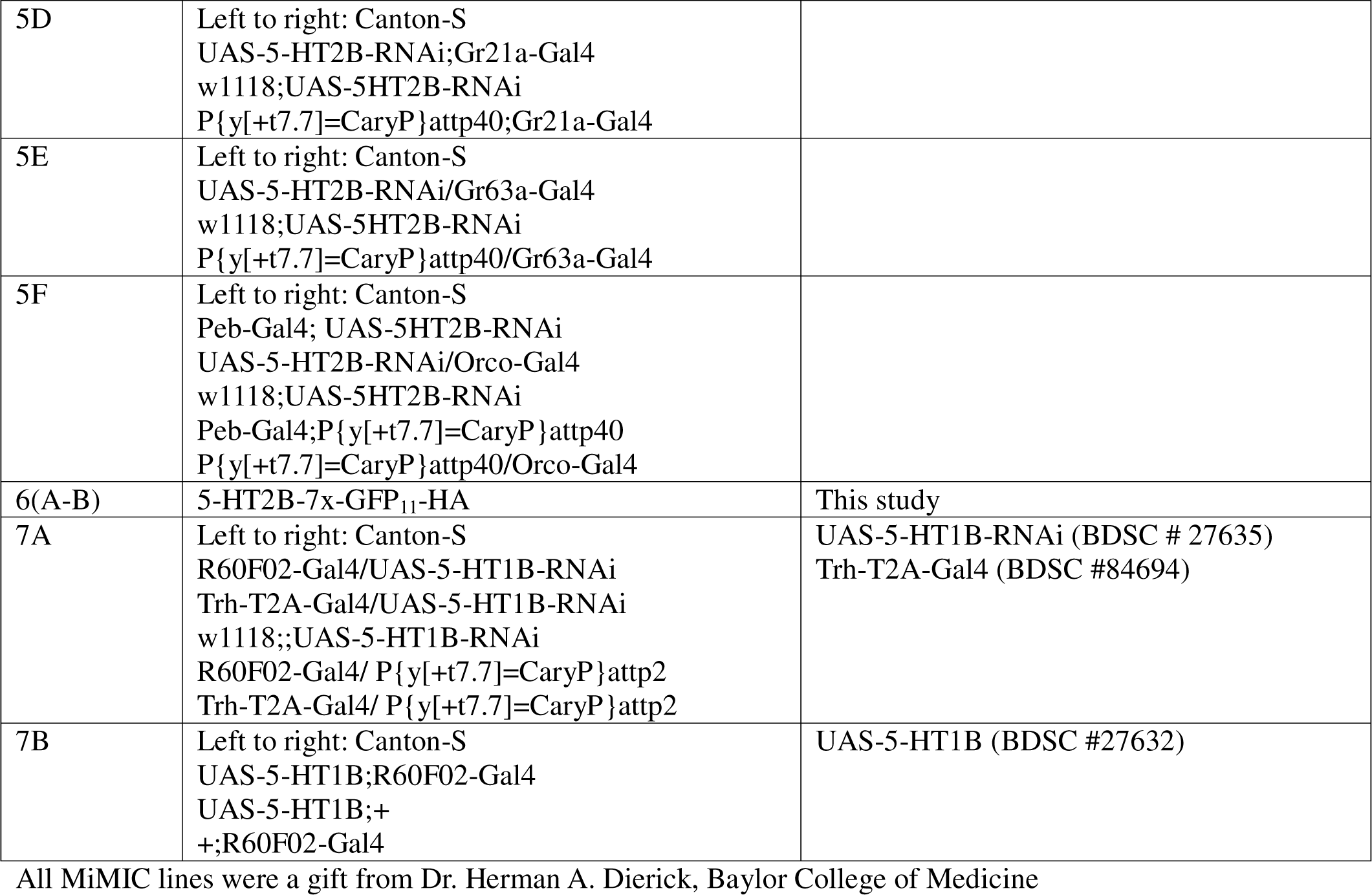
List of Flies used in this study.

#### Image Analysis

For measuring the volume of the glomerulus, we performed image segmentation by specifying the borders of the specific glomerulus across the Z-stack of the confocal image based on n-cad staining using the segmentation plugin in Image J. The volume of the selected glomeruli was obtained by 3-D analysis of the segmented image using the MorphoLibJ^43^ plugin in ImageJ. Absolute volume is computed in this plugin by multiplying the number of voxels comprising the selection with the volume of individual voxel. The resultant glomeruli volumes were found to be consistent with those previously reported^21,27^.

The fluorescence intensity measurements in Figure S2 were normalized for each glomerulus against the fluorescence intensity of the VL1 glomerulus. This was done to show the fold-difference in 5-HT2B expression in the selected glomeruli in comparison to a glomerulus of average 5-HT2B expression. The fluorescence intensity measurements reported in Figure 6B were obtained using previously described methodology in ImageJ^44^. The corrected total cell fluorescence for each sample reported was calculated by subtracting the product of the area of the AL and mean background fluorescence of the brain outside the AL from the integrated density of the AL.

Approximately 15 samples were analyzed for each genotype, age, or odor exposure unless otherwise noted. An unpaired Student’s t-test was used to compare the difference in glomerular volumes of air and CO_2_ exposed flies or fluorescence intensity between sexes or across fly of different ages. The p-values are indicated as * for p < 0.05 and as N.S for non-significance (p > 0.05) unless otherwise noted.

## RESULTS

### 1. Olfactory CPP requires the release of 5-HT by the serotonergic neurons

In *Drosophila*, the olfactory CPP manifests as a change in the volume of the glomerulus innervated by OSNs responsive to the odor used as a stimulus (Figures 1A-B).^20,21,27^ Consistent with previous reports,^20,21,27^ in flies where 5-HT transmission is intact, the V-glomerulus increased in volume in flies exposed to CO_2_ compared to air exposed (Figure 1B). In *Drosophila* and other holometabolous insects, a single pair of serotonergic neurons called the <u>C</u>ontralaterally projecting <u>S</u>erotonin immunoreactive <u>D</u>euterocerebral <u>n</u>eurons (CSDns) innervate the AL^47–49^ (Figure 1C) and supply serotonin.^48–51^ Previous work from our lab has shown that expression of the Vmat-RNAi transgene in the CSDns successfully eliminates 5-HT mediated responses in the AL.^41,50^ We thus assessed the role of 5-HT in CPP in the *Drosophila* olfactory system by measuring the structural plasticity induced in the CO_2_ sensitive V-glomerulus upon chronic exposure to 5% CO_2_. ^20,21,27^ We employed the R60F02-Gal4 promoter line that labels the CSDns^52^ to prevent serotonin release by knocking down Vmat in these cells via expression of Vmat RNAi. Chronic CO_2_ exposure to flies deprived of CSDn serotonin output failed to undergo structural plasticity in the V (Figure 1D), suggesting that serotonin release is required for structural plasticity.

**Figure 1.**
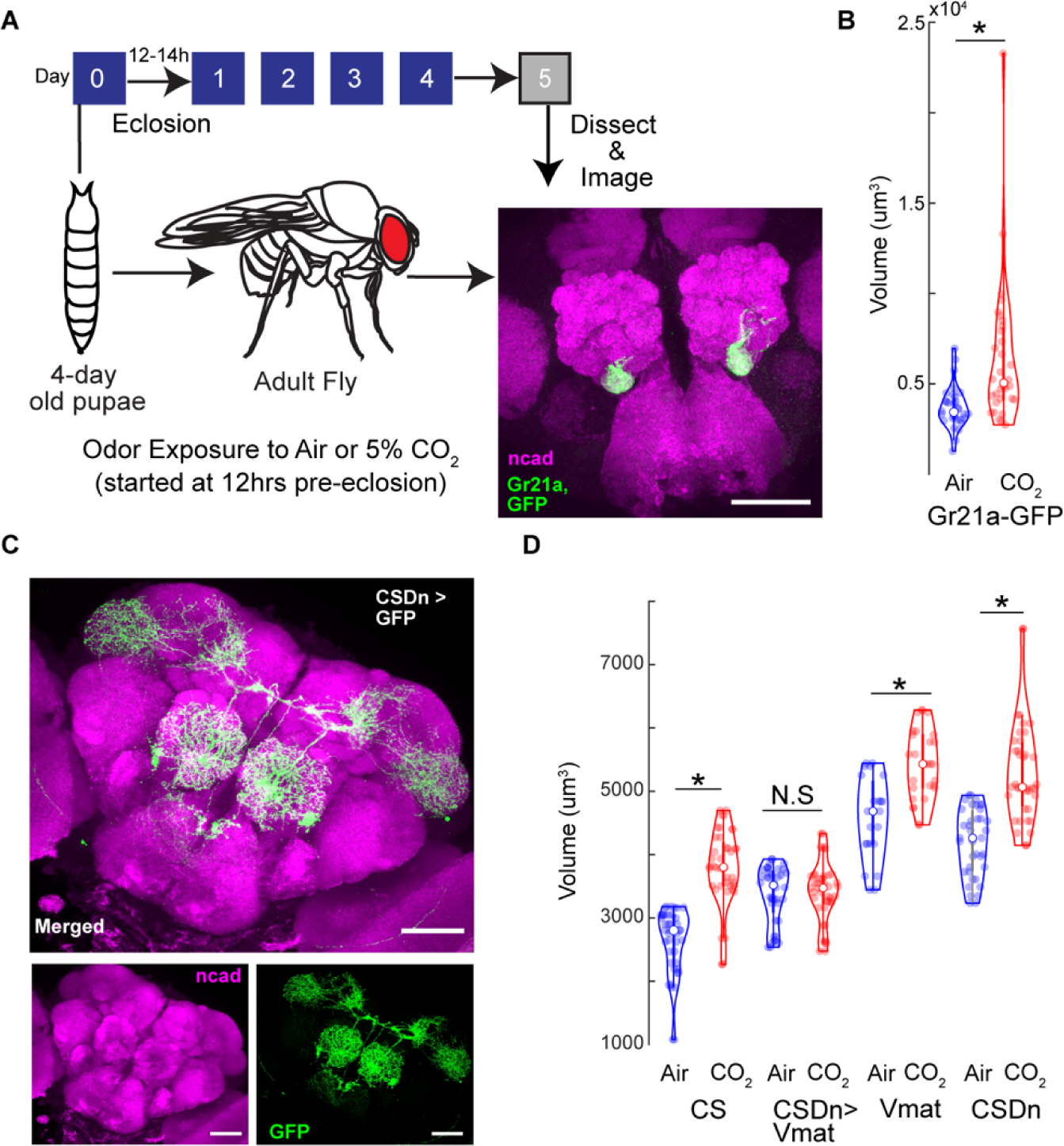
Blocking serotonin release from CSDns prevents structural plasticity during the critical period. (A) Schematic of the experimental protocol. 4-day old pupae are collected and subject to 5% CO_2_ for 5 days. On day 5 after eclosion, flies are collected and stained with n-cadherin and imaged under a confocal microscope to analyze structural plasticity. Confocal maximum intensity projection of AL by CO_2_ responsive V-glomerulus in green co-labeled for n-cadherin (magenta). (B) Quantification of V-glomerulus volumes comparing air (right) and 5% CO_2_ (left) exposed flies during the critical period. Genotype shown here is endogenously expressed GFP under the Gr21a promoter (Gr21aGFP). (C) Confocal maximum intensity projections of AL innervation by CSDns (CSDn-Gal4 > UAS-mcd8::GFP, green) co-labeled for n-cadherin (magenta). (D) Quantification of V glomerulus volumes comparing air and 5% CO_2_ exposed flies during the critical period. Four genotypes are shown here from left to right : Canton-S (wildtype), CSDn-Gal4>UAS-Vmat-RNAi (CSDn targeted Vmat knockdown), w1118;;UAS-Vmat-RNai (background control for Gal4) and y,v;; CSDn-Gal4 > RNAi background (background control for RNAi). * indicates p < 0.05; N.S indicates p > 0.05; n >=15 The scale bar indicates 50 μm in all cases

### 2. Serotonin modulates the olfactory CPP via multiple receptor targets

Having established that 5-HT plays a role in the olfactory CPP in *Drosophila*, we wished to determine the cellular and molecular targets by which 5-HT was exerting its impact. 5-HT mediates its effect in cells by concentration dependent activation of its cognate receptors. In *Drosophila*, there are 5 serotonin receptors (5-HTRs): 5-HT1AR, 5-HT1BR, 5-HT2AR, 5-HT2BR and 5-HT7R.^53–56^ We systematically interrogated 5-HTR signaling to determine which receptors are involved in mediating structural plasticity during the critical period (Figure 2). For this, we used the same experimental paradigm as before and employed the null mutants of the 5-HTRs generated by a CRISPR knock in strategy to replace all or parts of the gene encoding the 5-HTRs with the GAL4 gene.^56^ Chronic exposure of CO_2_ in flies with mutations in the 5-HT1BR and 5-HT7R (Figures 2C&F) failed to demonstrate structural plasticity in the V-glomerulus during the critical period. Since the 5-HT2BR homozygous mutants were not viable in our hands, we employed a slightly modified strategy to investigate its effects on CPP. We crossed the heterozygous 5-HT2BR mutants expressing Gal4 under the 5-HT2BR promoter to induce expression of 5-HT2B-RNAi. This 5-HT2BR deficient state was sufficient to block structural plasticity in the V-glomerulus during the critical period (Figure 2E). In contrast, we still observed structural plasticity in 5-HT1AR and 5-HT2AR mutants upon chronic CO_2_ exposure during the CP (Figures 2B&D) indicating that these two receptors do not underlie the effects of 5-HT on early life olfactory plasticity. Together, these results indicate that 5-HT1BR, 5-HT2BR and 5-HT7R are required during the critical period.

**Figure 2.**
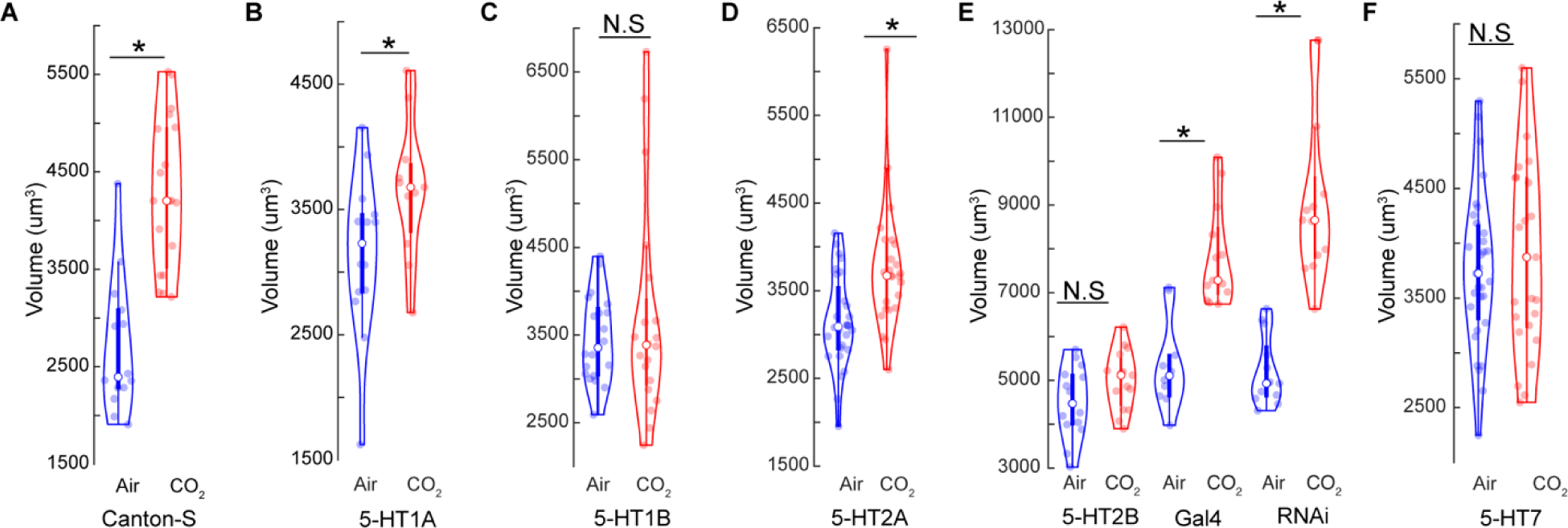
Serotonin acts through multiple receptors during the critical period. (A) Comparison of V-glomerulus volumes in air and 5% CO_2_ exposed brains in wildtype Canton-S flies. (B-D, F) Comparison of V-glomerulus volumes in air and 5% CO_2_ exposed brains of 5-HT1AR (B), 5-HT1BR (C), 5-HT2AR (D) and 5-HT7R (F) knockout flies. (E) Comparison of V-glomerulus volumes in air and 5% CO_2_ exposed brains of flies with genotypes (left to right): 5-HT2B knockdown in 5-HT2B heterozygous mutants, UAS-5HT2B-RNAi in Gal4 knock in background, heterozygous 5-HT2B mutant with Gal4 knock in RNAi background (y,v;5HT2B[Gal4] > TRiP background). * indicates p < 0.05; N.S indicates p > 0.05; n >= 15 in all cases

Previous studies demonstrated that all five 5-HT receptors are expressed by distinct neuron types in the AL.^57^ These studies relied on GFP expression induced by the Gal4 protein expressed from the endogenous promoter. While this approach reveals which cells classes express which 5-HTRs, the method does not easily identify where and when the receptors are trafficked within the neurons. We therefore generated flies with an endogenous HA-tag and the GFP_11_ fragment on 5-HTRs.^58–60^ We employed a CRISPR-Cas9 based strategy^39,40^ to generate these flies. The split-GFP is an elegant tool that consists of splitting the superfolder GFP (sfGFP) between the beta-strand 10 and 11 to generate two non-fluorescing, self-complementing fragments: GFP_1-10_ and GFP_11_.^61,62^ Only cells that would simultaneously express both fragments will be able to form the complete sfGFP molecule that would fluoresce (Figure S1A). Additionally, the HA tagged 5-HTRs would enable us to locate 5-HTR expression in all cells by immunolabeling against HA (Figure S1B). Together, this strategy allows us to simultaneously visualize the localization of 5-HTRs and determine the cell types that express them (Figure S1C-F). We found that the expression patterns of these 5-HTR lines labeled using the MiMIC-5HTR-Gal4 drivers to be consistent with previously reported expression patterns of the 5-HTRs^45,57^ using more traditional GAL4/UAS approaches (Figures 3A-E).

**Figure 3.**
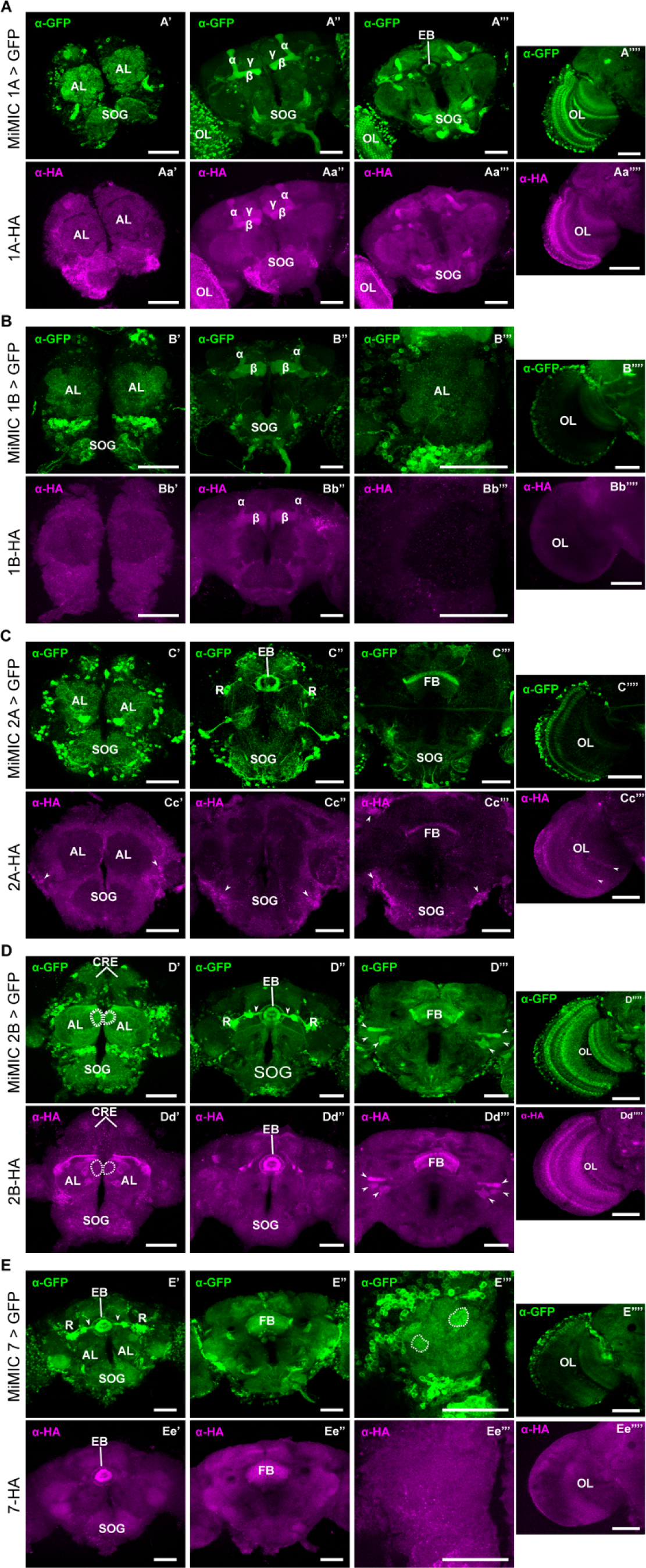
Distribution of 5-HTRs in adult *D. melanogaster* brains. (A) Brain sections indicate 5-HT1A expression in the AL, suboesophageal ganglion (SOG), MB, ellipsoid body (EB), and optic lobe (OL). As shown in (A’’) and (Aa’’), both α-HA and α-GFP indicate expression in all three lobes of MB, including α, β, and γ lobes. Note, as shown in (Aa’’’), 5-HT1A expression in EB is largely devoid revealed by α-HA, contrary to the α-GFP staining pattern shown in (A’’’). (B) Brain sections indicate 5-HT1B expression in SOG, AL, MB and OL. In contrast to the MiMIC approach, which labels cell bodies broadly in the brain, immunostaining against the HA tag show significant distribution of the receptor only in the α and β lobes of MB. (C) Brain sections indicate 5-HT2A expression in many areas such as AL, SOG, EB, the fan-shaped body (FB), and optical lobe (OL). Among all these areas the staining by α-HA shows weak signals in FB (Cc’’’) and probably in OL (Cc’’’’, indicated by the arrow heads). The contrast of the images from the α-HA channel were elevated to see weak signals, resulting in irregular signal presentation which might be artefacts, as indicated by the arrow heads in (Cc’-Cc’’’). (D) Brain sections indicate 5-HT2B expression in AL, crepines (CRE), SOG, EB, FB, and OL. In ALs, the MiMIC approach indicates a relatively high expression in a medial glomerulus, as emphasized by the white dashed circle, which is not consistent in the α-HA channel (Dd’). It is noticeable that localization of this receptor in the R neurons (R) and their arborizations toward EB (D’’, arrow heads) is missing in the staining pattern revealed by α-HA (Dd’’). Arrow heads in (D’’’) and (Dd’’’) indicate some unknown structures stained in both channels. (E) Brain sections indicate 5-HT7 expression in SOG, EB, FB, and (E’’’’/Ee’’’’) OL. In the antennal lobe labeled with the MiMIC approach, extraordinary signals were observed in an anterior dorsal glomerulus as indicated by dashed circle on the right side, and a posterior lateral glomerulus as indicated by dashed circle on the left side (E’’’). In contrast, no prominent signals showed in these two glomeruli as indicated by the α-HA approach (Ee’’’). Signals were observed in the optical lobe (OL) revealed by both approaches (E’’’’/Ee’’’’). For both (E’’’/Ee’’’) and (E’’’’/Ee’’’’), the brain was oriented with the lateral toward left and the dorsal upward. The scale bar indicates 50 μm in all cases.

Next, we investigated the differential expression patterns of the 5-HTRs in the olfactory processing centers of the brain. The 5-HT1Rs are expressed in varying degrees within the AL and MBs (Figure 3A). The 5-HT2ARs are mostly expressed in the cells surrounding and innervating the AL, most likely the LNs and PNs (Figures 3C’ and Cc’). Remarkably 5-HT2BRs is not uniformly expressed throughout the AL (Figures 3D’,Dd’ and S2A-I). This indicates varying levels of 5-HT2BR mediated serotonergic modulation in the OSNs. Consistent with prior reports,^57^ we found most of the 5-HT2BR expression in the AL to be in the OSNs. When we removed the antennae or the maxillary palps that houses the cell body and dendrites of OSNs, 5-HT2BR expression in the related AL region the OSNs project to is eliminated (Figure S2J). Similarly, we were able to selectively knockdown 5-HT2BRs expression using Gal4 drivers for OSN subtypes using an RNAi against 5-HT2BR (Figures S2K,L). The AL neuropil is innervated by various cells including OSNs, PNs, LNs and CSDns. We found most of the 5-HT7R expression in the cells surrounding the AL, most likely in the LNs and PNs (Figures 3E’,E’’’). We also found unusually high GFP labeling in two glomeruli in the AL (Figure 3E’’’) but no corresponding HA labeling (Figure 3Ee’’’) for 5HT7R expression using the MiMIC-5HT7R-Gal4 promoter line. Taken together, these results show that 5-HT targets multiple components of olfactory processing through distinct receptors. Therefore, serotonin could target multiple serotonergic receptors on distinct cell types to mediate their effects during the olfactory critical period.

### 3. Serotonin modulates distinct components of sensory processing during the critical period

Next, we wanted to isolate the neuronal basis of 5-HTR signaling that modulates CPP. Earlier studies have identified a crucial role of inhibitory, GABAergic LNs, namely LN1 and LN2 during the olfactory critical period.^20,21,27^ Previous work in our lab has identified a distinct population of 5-HT7R expressing GABAergic LNs (R70A09-Gal4) that are responsive to low 5-HT concentrations and modify odor coding in the AL.^41^ As a population, these LNs innervate all glomeruli including the V-glomerulus (Figure 4A). We also found that R70A09-GAL4 LNs express 5-HT7Rs (Figure 4B) and show almost a complete overlap with LN1 neurons (Figures 4C,D) and a partial overlap with the LN2 neurons (Figures 4E,F). The LN1 neurons have been previously implicated to induce an increase in the number of PN arbors leading to structural plasticity during the critical period.^20,21^ We therefore sought to determine if these LNs are the target for 5-HT modulation via the 5-HT7Rs and found that knocking down 5-HT7Rs in the R70A09 LNs was sufficient to prevent CPP in the V-glomerulus (Figure 4G). In *Drosophila*, 5-HT7Rs are known to activate an adenylate cyclase that results in an increase in cytosolic cAMP.^63,64^ When we overexpress the cAMP specific phosphodiesterase *dunce* to deplete cAMP selectively in the R70A09 LNs keeping the 5-HT7Rs and adenylate cyclase intact, CPP in the V-glomerulus is abolished (Figure 3H). These results indicate that both 5-HT7R signaling and cAMP in R70A09-GAL4 LNs play an important role during the CP that ultimately permit the induction of structural plasticity in the cognate glomerulus.

**Figure 4.**
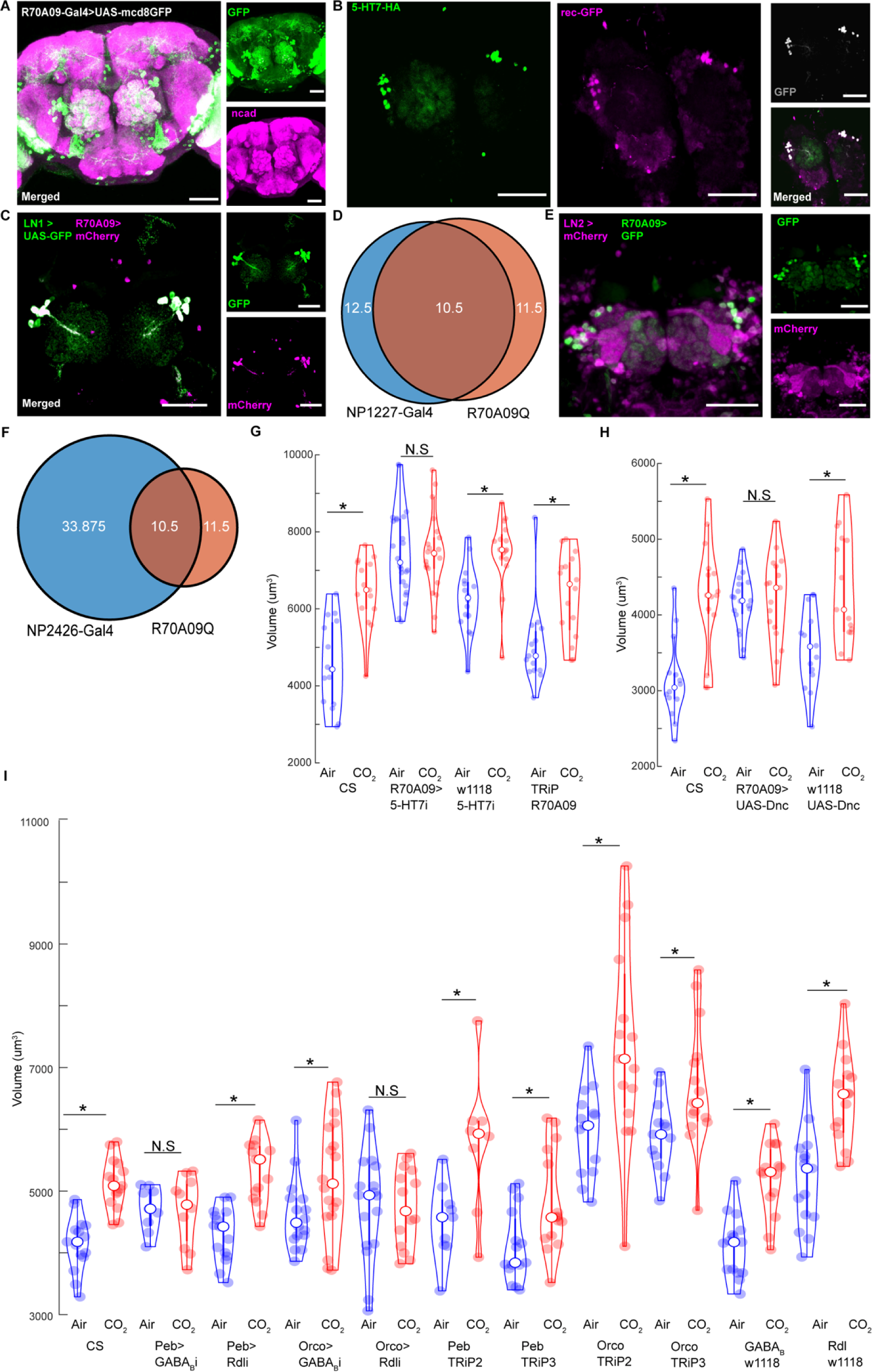
5-HT7 targets GABAergic inhibition in the primary olfactory circuit. (A) Confocal maximum intensity projection of the R70A09 LNs expressing GFP in green, co-labeled for n-cadherin in magenta. The V-glomerulus is circled in the merged and GFP channels. (B) Confocal maximum intensity projection of the AL showing R70A09 LNs co-labeled for 5-HT7-HA in green, reconstituted GFP in magenta, GFP in grey and the merged channel. (C) Confocal maximum intensity projection of the AL showing the overlap between R70A09 LNs in magenta and LN1 neurons in green. (D) The R70A09 line labels 11-12 LNs, LN1 labels 12-15 LNs per hemisphere. There is a total overlap of 10-11 cells between R70A09 and LN1 population per hemisphere. (E) Confocal maximum intensity projection of the AL showing the overlap between R70A09 LNs in green and LN2 in magenta. (F) The R70A09 line labels 11-12 LNs, LN2 labels 33-44 LNs per hemisphere. There is a total overlap of 10-11 cells between R70A09 and either LN1 or LN2 population per hemisphere. (G) Quantification of V glomerulus volumes comparing air and 5% CO_2_ exposed flies during the critical period. Four genotypes are shown here from left to right: CS (Canton-S wildtype), R70A09>5-HT7i (R70A09 targeted 5-HT7 knockdown), w1118 5-HT7i (background control for Gal4) and TRiP R70A09 (background control for RNAi). (H) Comparison of V-glomerulus volumes in air and 5% CO_2_ exposed brains of flies with overexpression of *dunce* in the R70A09 LNs. 3 genotypes are shown here from left to right: CS (Canton-S wildtype), R70A09>UAS-Dnc (*dunce* overexpression in R70A09 LNs), w1118 UAS-Dnc (background for Dnc and R70A09). (I) Comparison of V-glomerulus volumes in air and 5% CO_2_ exposed brains of flies with GABA receptor knockdown in OSNs. 11 genotypes are shown here from left to right: CS (Canton-S wildtype), Peb>GABA_B_i (GABA_B_ knockdown in all OSNs), Peb>Rdli (GABA_A_ knockdown in all OSNs), Orco>GABA_B_i (GABA_B_ knockdown in Or83b OSNs), Orco>Rdli (GABA_A_ knockdown in Or83b OSNs), Peb TRiP2 (background control for Rdl RNAi crossed with Peb-Gal4), Peb TRiP3 (background control for GABA_B_ RNAi crossed with Peb-Gal4), Orco TRiP2 (background control for Rdl RNAi crossed with Orco-Gal4), Peb TRiP3 (background control for GABA_B_ RNAi crossed with Orco-Gal4), w1118 GABA_B_i (background for Gal4 crossed with GABA_B_-RNAi), w1118 Rdli (background for Gal4 crossed with Rdl-RNAi). * indicates p < 0.05; N.S indicates p > 0.05. n>=15. The scale bar indicates 50 μm in all cases.

The R70A09 LNs that express 5-HT7 receptors are GABAergic in nature and release GABA upon activation.^41^ The pan-glomerular innervation of these LNs implies that they release GABA all over the AL. Apart from these LNs, other GABAergic LNs also exist in the AL that can be sensitive to serotonergic modulation. Previous work has shown that GABA released from the GH298 LNs which are distinct from the R70A09 LNs mediate glomerulus selective presynaptic divisive gain control in Or83b expressing OSNs in adult *Drosophila* but do not affect CO_2_ responses in the Gr21a expressing OSNs.^65^ Consistent with these studies, knocking down GABA_B_ and GABA_A_ receptors respectively in the CO_2_ sensing OSNs and PNs was not sufficient to block structural plasticity during the CP.^20,23^ However, 5-HT7R mediated activation in the R70A09 LNs and thereby GABA release is important for inducing CPP. Therefore, it is likely that R70A09 LN activation linked GABA release during the critical period could lead to network level changes in the AL that ultimately facilitate structural plasticity in the cognate glomerulus. We asked if the two GABA receptors expressed in the fly, the GABA_A_ and GABA_B_ receptors are required for global inhibition in the OSNs during the critical period. When we knock down GABA_B_ receptors in all OSNs we saw no structural plasticity in the V-glomerulus (Figure 4I). In contrast, knocking down GABA_A_ receptors in all OSNs did not hinder CPP in the V (Figure 4I). Finally, we targeted Or83b OSNs in the AL using the Orco-Gal4 promoter line. This enabled targeting multiple OSNs responsive to different odors^41,65^ but not the CO_2_ sensing ones. Surprisingly, flies expressing GABA_A_ RNAi in Or83b OSNs failed to undergo structural plasticity in the V upon CO_2_ exposure (Figure 4H). However, knocking down GABA_B_ in the Or83b OSNs was not sufficient to prevent structural plasticity in the V in response to CO_2_ (Figure 4I). This shows that GABAergic inhibition during the critical period is required in a broader sub-population of OSNs in the AL but not selectively in the cognate glomerulus. In fact, GABA targets distinct OSNs through GABA_A_ and GABA_B_ receptors to facilitate CPP in the V glomerulus.

Next, we asked which cells projecting to the AL could be the source of 5-HT2BR mediated serotonergic modulation of the olfactory CPP. We hypothesized that 5-HT targets 5-HT2BRs on OSNs during the critical period. Previous research has shown that the 5-HT2BRs are expressed by all OSNs and a few LNs and PNs in the AL.^57^ We showed earlier in Figure S2J that the majority of the 5-HT2BR expression in the antennal lobe is due to their expression by the OSNs. Therefore, we selectively knocked down expression of the 5-HT2BRs in the V-glomerulus OSNs (Figure 5). In the CO_2_ detecting OSNs, two chemoreceptors, Gr21a and Gr63a are co-expressed to form a functional CO_2_ responsive odor receptor^66^. However, in our 5-HT2BR knockdown experiments, we observed residual 5-HT2BR expression in the V-glomerulus using the Gr21a-GAL4 driver line (Figure 5A). Therefore, 5-HT2BR knockdown driven by the Gr21a-Gal4 line was not sufficient to prevent CPP in the V-glomerulus (Figure 5D). In contrast, driving the 5-HT2BR RNAi using Gr63a-Gal4 significantly reduced 5-HT2BR expression in the V-glomerulus without impacting expression in the rest of the AL (Figure 5B). This selective knockdown of the 5-HT2BRs by the Gr63a-Gal4 line was sufficient to prevent the induction of structural plasticity in the V-glomerulus (Figure 5E) suggesting that 5-HT2BR expression is required by OSNs within the glomerulus expanding during the CP. To determine if 5-HT2BR expression by OSNs in other glomeruli is required for CPP in the V-glomerulus, we next extended the 5-HT2BR knockdown using drivers expressed broadly in all OSNs (Peb-Gal4) or in many OSNs except those projecting to the V and a few other glomeruli (Orco-Gal4). We found that flies that expressed 5-HT2B RNAi in all OSNs (Figure 5C), failed to undergo structural plasticity in the V-glomerulus in response to chronic CO_2_ exposure (Figure Figure 5F). In contrast, 5-HT2B knockdown in multiple OSNs using the Orco-Gal4 driver line did not show any deficits in structural plasticity in response to CO_2_ (Figure 5F). Together, these results show that the 5-HT2BR expression is required in cognate ORNs of the V glomerulus for proper expression of CPP.

**Figure 5.**
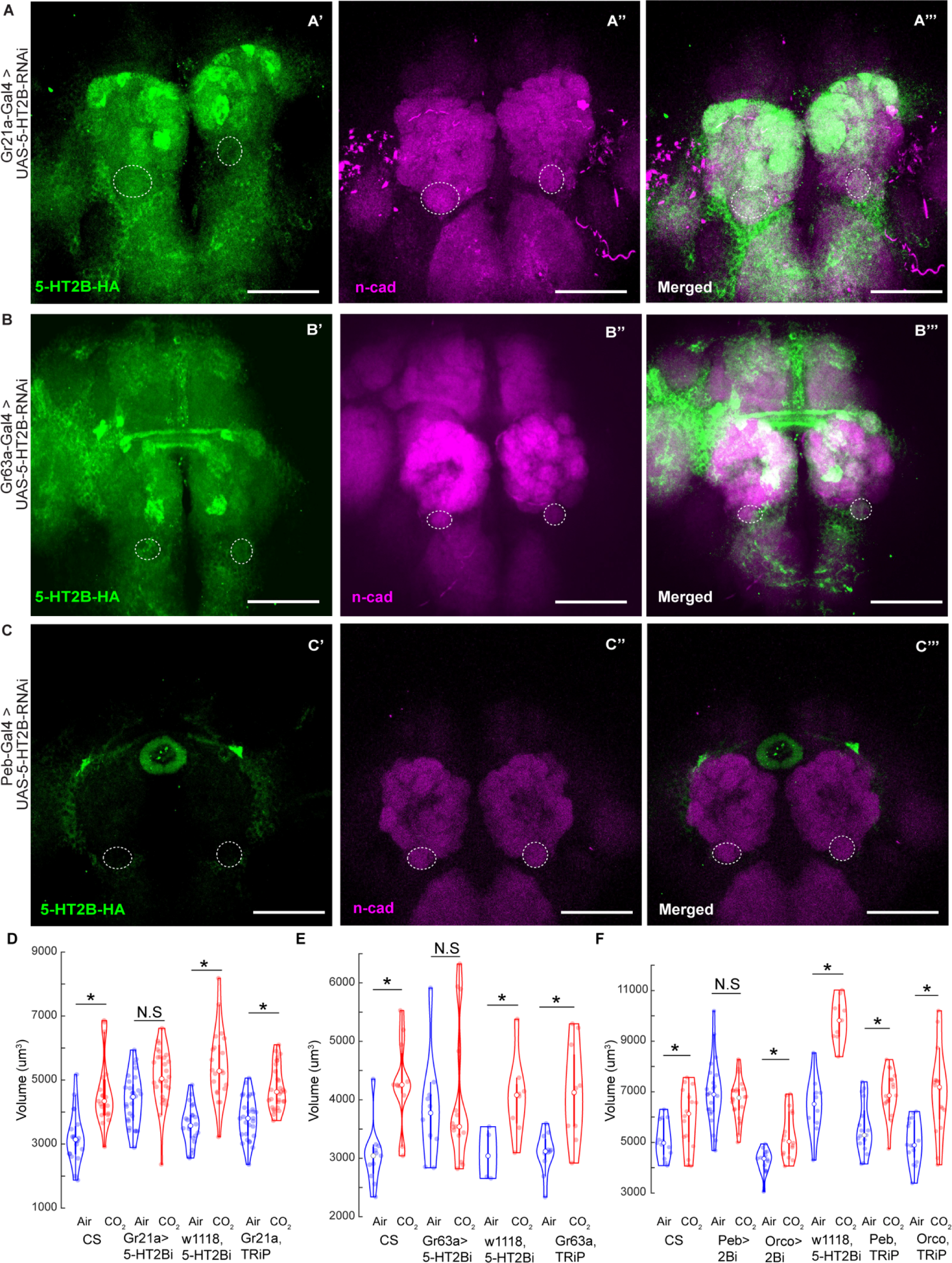
5-HT2BRs are required in OSNs during the critical period. (A) Insufficient 5-HT2B knockdown in the CO_2_ sensing OSNs by Gr21a-Gal4. The V glomerulus is circled on all three channels. (B) 5-HT2B knockdown in the CO_2_ sensing OSNs by Gr63a-Gal4. The V glomerulus is circled on all three channels. (C) 5-HT2B knockdown in all OSNs but no other brain regions by Peb-Gal4. The V glomerulus is circled on the n-cad (magenta) and merged channels. (D) Quantification of V glomerulus volumes comparing air and 5% CO_2_ exposed flies during the critical period. Four genotypes are shown here from left to right: CS (Canton-S wildtype), Gr21a>5-HT2Bi (5-HT2B knockdown in CO_2_ OSNs), w1118 5-HT2Bi (background control for Gal4) and Gr21a, TRiP (background control for RNAi). (E) Quantification of V glomerulus volumes comparing air and 5% CO_2_ exposed flies during the critical period. Four genotypes are shown here from left to right: CS (Canton-S wildtype), Gr63a>5-HT2Bi (5-HT2B knockdown in CO_2_ OSNs), w1118, 5-HT2Bi (background control for Gal4) and Gr63a, TRiP (background control for RNAi). (F) Quantification of V glomerulus volumes comparing air and 5% CO_2_ exposed flies during the critical period. Six genotypes are shown here from left to right: CS (Canton-S wildtype), Peb>5-HT2Bi (5-HT2B knockdown in all OSNs), Orco> 2Bi (5-HT2B knockdown in Or83b OSNs), w1118,5-HT2Bi (background control for Gal4), Peb, TRiP (background control for 5-HT2B-RNAi) and Orco,TRiP (background control for 5-HT2B-RNAi). * indicates p < 0.05; N.S indicates p > 0.05. n>=15. The scale bar indicates 50 μm in all cases.

We also observed glomerulus specific differences in the expression levels of the 5-HT2BRs. (Figure S2A-I). Therefore, to determine if expression of the 5-HT2BRs within the AL varies post-eclosion, we employed the 5-HT2BR-HA tagged recombinant flies. We found that the expression of the 5-HT2BRs increases significantly post-eclosion and reaches its peak at day 2 or 48-hour post eclosion (Figures 6A,B), which coincides with the closing of the critical period.^21,27^ After day 2, the 5-HT2BR expression does not vary significantly as we saw no difference in the corrected total fluorescence between day 2, day 4 and day 5 post eclosion. Taken together, these results show that 5-HT targets both excitatory and inhibitory neurons within the antennal lobe via distinct receptors.

**Figure 6.**
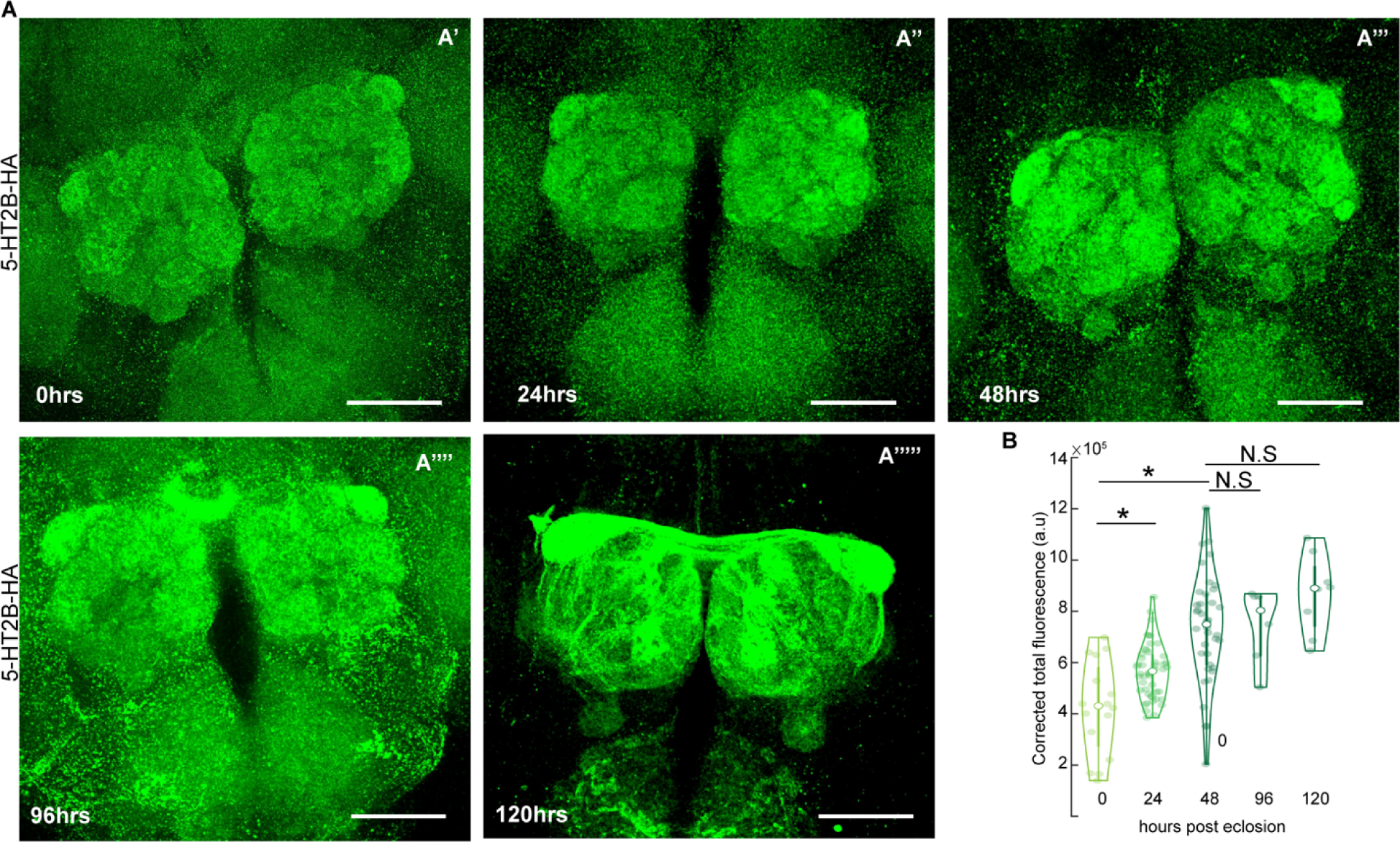
5-HT2BR expression in the AL varies during the critical period. (A) 5-HT2B expression levels in the AL as indicated by HA staining in green at different life stages of the fly post eclosion: 0 hrs or freshly eclosed (A’) 24hrs (A’’), 48hrs (A’’’), 96hrs (A’’’’) and 120hrs (A’’’’’). (B) Total corrected cell fluorescence indicating 5-HT2B expression levels in the AL of the fly at different ages from left to right: 0hrs or freshly eclosed, 1 day or 24 hrs post eclosion (p.e.), 2 days or 48hrs p.e., 4 days or 96hrs p.e., and 5 days or 120 hrs p.e. ** indicates p < 0.05; N.S indicates p > 0.05. The scale bar indicates 50 μm in all cases. n >= 15.

#### Autoregulation of serotonergic neurons during the critical period

Finally, we sought to determine the neurons for whom expression of the 5-HT1BR is required for the olfactory CPP. Within serotonergic neurons, the 5-HT1BRs often act as auto receptors by either inhibiting the release of 5-HT^67–70^ or by modulating serotonin reuptake by upregulating SERT activity and clearance rate.^71^ Therefore, we knocked down 5-HT1B receptors in the CSDns, release of 5-HT from which is required during critical period (Figure 7A). We found that 5-HT1B signaling in CSDns is required for glomerular specific volume increase during the critical period. Similarly, knocking down 5-HT1BRs in all serotonergic neurons in the brain using a Gal4 promoter line Trh-Gal4, was able to block CPP in the V-glomerulus (Figure 7A). The CSDns release 5-HT upon activation. Since the 5-HT1BRs are inhibitory in nature, activation of 5-HT1BRs on CSDNs will inhibit 5-HT release from them. We also know that release of 5-HT from the CSDns is important during the critical period. Therefore, if we overexpress 5-HT1BRs in the CSDns there will be a stronger inhibition in the CSDns most likely preventing CPP. Consistent with our hypothesis, we saw that 5-HT1BR overexpression on CSDNs prevents CPP in the V-glomerulus following CO_2_ exposure (Figure 7B). Together these results indicate that 5-HT levels need to be tightly controlled to induce CPP.

**Figure 7.**
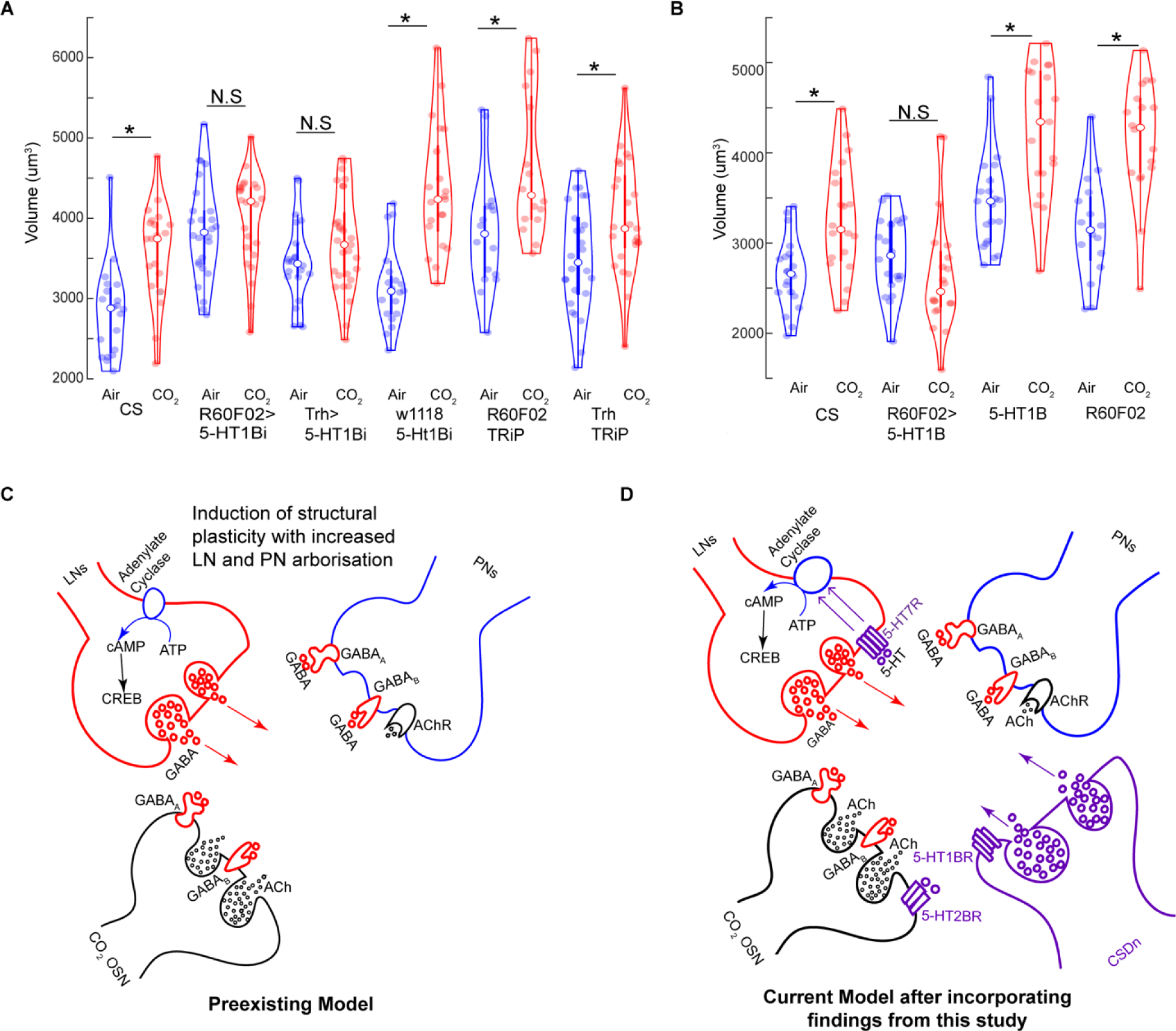
Serotonergic signaling during the critical period. (A) Quantification of V glomerulus volumes comparing air and 5% CO_2_ exposed flies during the critical period. Six genotypes are shown here from left to right: CS (Canton-S wildtype), R60F02>5-HT1Bi (RNAi knockdown in CSDns), Trh>1Bi (5-HT1B knockdown in all serotonergic cells), w1118,5-HT1Bi (background control for Gal4), R60F02, TRiP (background control for 5-HT1B-RNAi crossed with CSDn line) and Trh,TRiP (background control for 5-HT1B-RNAi crossed with Trh line). (B) Quantification of V glomerulus volumes comparing air and 5% CO_2_ exposed flies during the critical period. Four genotypes are shown here from left to right: CS (Canton-S wildtype), R60F02>5-HT1B (Overexpression of 5-HT1B in CSDns), 5-HT1B (control for UAS), R60F02 (control for Gal4). (C) During the critical period, chronic odor exposure leads to OSN dependent activation of LNs and PNs. Activation of GABAergic LNs induces GABA release that modulates OSN and PN responses. cAMP dependent mechanisms in GABAergic LNs lead to CREB dependent gene transcription that promotes to structural plasticity in the LN and PN arbors resulting in glomerulus specific volume increase. (D) Our results (in pink) show 5-HT also plays a role within the existing model of CPP. 5-HT is released from the CSDNs and is tightly regulated by 5-HT1BRs during the critical period. Differential expression of 5-HT2B neurons on the OSNs regulates structural plasticity in the LNs and PNs. 5-HT7 mediated GABAergic LN activation interacts with the preexisting model of cAMP dependent gene transcription to facilitate CPP. GABAergic signaling from the LNs modulate global OSN activation levels during the critical period. * indicates p < 0.05; N.S indicates p > 0.05. n>=15

## DISCUSSION

There are many cellular and molecular components that contribute to CPP. Serotonin is elegantly positioned to affect CPP because a diverse set of 5-HT receptors are broadly expressed throughout the network. The genetically accessible olfactory circuit of *Drosophila* allowed us to isolate the effects of serotonergic modulation in the critical period exclusively within the olfactory circuit via regulating serotonin release from the CSDns and selectively knocking down 5-HTRs in specific cell types within the olfactory circuit. Our results indicate that 5-HT modulates both excitatory and inhibitory elements in the olfactory circuit during the critical period.

### 5-HT modulates inhibitory LN circuits that underlie critical period plasticity

Critical periods are known to be tightly regulated by the maturation of inhibitory circuits. In fact, the emergence and maturation of GABAergic inhibitory local interneurons (LNs) in mammals^30,72^ are known to improve the signal to noise ratio by improving their excitation/inhibition balance^30^ in visual^73^, auditory^29^, and somatosensory^74^ cortices during the critical period. Previous investigations of the olfactory critical period in *Drosophila* have identified a key role of two distinct inhibitory, GABAergic LN populations (LN1 and LN2) in modulating PN output and structural plasticity upon chronic odor exposure during the critical period.^20,27^ The LN1 sub-population labelled by the NP1227 Gal4 line showed small, statistically insignificant increments in its dose-response curve during chronic CO_2_ exposure. In contrast, the LN2 LNs labelled by the NP2426-Gal4 line showed significant increases in its cytosolic Ca^2+^ upon chronic CO_2_ exposure during the critical period.^27^ We identified that 5-HT7R mediated serotonergic modulation within this circuit activates cAMP dependent mechanisms of gene expression in LN1 neurons that then induces the volume changes.

The CSDns maintain reciprocal connections with the inhibitory, GABAergic, and glutamatergic LNs within the AL.^48^ These LNs are critical for network level inhibition in the AL as they tone down PN output before it reaches the MB and LH.^65,75–80^ Following odor exposure, CSDn inhibits some LN types via 5-HT. In turn, the CSDns are inhibited by both GABAergic and glutamatergic inhibition.^50^ Thus, the CSDns could modulate network level inhibition in the AL via serotonergic modulation and can themselves undergo inhibition based on the network wide inhibitory dynamics established by the LNs. While blocking 5-HT release from the CSDns prevented the availability of synaptic 5-HT and thereby CPP, the AL neurons also had access to basal 5-HT levels released by the remaining 108 serotonergic neurons in *Drosophila*. A prime candidate of basal 5-HT modulation is the 5-HT7R expressing R70A09 GABAergic LNs we identified to be required during the critical period.^41^ These R70A09 LNs mediate subtractive gain control in the PNs and thereby downregulate global PN responses^41^. Our results indicate that although the basal 5-HT levels are trivial during the critical period, 5-HT7R mediated modulation of R70A09 is required during the critical period. This implies that these cells are most likely playing a crucial role during the critical period in maintaining the inhibitory tone in the AL that is conducive to structural plasticity during the critical period.

In *Drosophila*, the Ca^2+^/calmodulin sensitive adenylate cyclase rutabaga (*rut*) acts as a coincidence detector for cytosolic Ca^2+^ increase and GPCR activation^20,81^ and converts ATP to cAMP. Rescuing (*rut*) in LN1 or GABA expressing glutamic acid decarboxylase (GAD-1) positive neurons in *rut^2080^* mutants was sufficient to reinstate CPP in those flies^20,21^. We found almost a complete overlap between 5-HT7 expressing R70A09 LNs and the LN1 neurons. Therefore, we can presume a serotonergic modulation in LN1 and consequently in the R70A09 LNs to be acting via 5-HT7 mediated cAMP increase that results in CREB dependent gene transcription. Our results show that depleting cAMP while keeping the 5-HT7Rs intact in the R70A09 LNs was sufficient to block CPP. The known organisms across phyla that express 5-HT7 all employ an adenylate cyclase dependent mechanism to increase cytosolic cAMP.^63,64,82–94^ Within the LN1 neurons, the adenylate cyclase *rutabaga* is required for CPP.^20,21^ It is likely that 5-HT7Rs expressed in these LNs modulate cAMP dependent gene transcription to facilitate structural plasticity during the critical period. This is also consistent with the known mechanism of 5-HT7R activity which increases intracellular cAMP levels in *Drosophila*.^64^ Future work is required to identify if 5-HT7Rs in *Drosophila* acts through the adenylate cyclase *rutabaga* to induce CREB dependent structural plasticity in the R70A09/LN1 neurons.

### 5-HT directly modulates excitatory OSNs during the critical period

In addition to impacting local interactions, serotonin also directly impacts 5-HT2BRs on OSNs during the critical period suggesting that there is direct modulation of primary sensory afferents by 5-HT. The differential expression levels exhibited by 5-HT2BRs following eclosion and until the end of the critical period (2 days post eclosion) in the AL is reminiscent of the patchy temporal expression of 5-HT2CRs in the kitten striatal cortex during the visual critical period.^33^ Since the ORNs are the primary source of 5-HT2BR expression in the AL, it is likely that the 5-HT2BR mediated serotonergic modulation adapts specifically to the odor environment presented to the fly during the critical period. This explains why knocking down 5-HT2BRs in the CO_2_ responsive OSNs prevents CPP in its cognate V-glomerulus. These results indicate that lower levels of 5-HT2BR expression are permissive to CPP while higher levels of 5-HT2BR signaling beyond the critical levels achieved at day 2 prevent CPP. Additionally, we found differential, patchy expression patterns of the 5-HT2BRs in distinct glomeruli of the antennal lobe during the critical period and in adults, which could also indicate odor dependent differential serotonergic modulation within distinct glomerulus in the AL. Future work directly correlating 5-HT2BR expression in individual glomerulus with the induction of CPP can shed light on the exact levels of 5-HT2BR modulation required during the critical period. Since the CO_2_ sensing OSNs do not undergo structural plasticity during the critical period^27^, downstream pathways by which 5-HT2BRs modulate OSNs during the critical period might shed light on how they indirectly affect structural plasticity in the LNs and PNs. The most likely mechanism could be via NMDA receptor dependent coincident detection that plays an important role in mediating glomerulus specific volume increase during the critical period.^20,21,27^ Thus, 5-HT can differentially modulate distinct glomeruli based on their 5-HT2BR expression levels upon a specific odor encounter during the critical period.

### Maintenance of optimum serotonin levels during the critical period

The balance of excitation and inhibition (E/I) within a network is fundamentally important for facilitating critical period plasticity.^95,96^ Intracellular electrophysiological recordings in mice indicate 5-HT acts by differentially modulating relevant excitatory and inhibitory synapses during the critical period.^97^ This indicates that 5-HT levels can play an important role in controlling permissive levels of excitation and inhibition during the critical period. Therefore, 5-HT levels and thereby 5-HT release permissive to CPP needs to be tightly controlled. Serotonin neurons are known to be modulated by 5-HT itself via expression of 5-HTRs.^64^ We show direct evidence of how serotonergic neurons modulate their own release to achieve optimum 5-HT levels through autoregulation. In the larval *Drosophila* nociceptive circuit, the inhibitory 5-HT1BRs expressed in a pair of serotonergic neurons directly inhibit sensory afferents and facilitate a form of experience dependent plasticity^98^. Similarly, the 5-HT1BRs are expressed by the CSDns^64^ and knocking down (Figure 7A) or overexpressing the 5-HT1BR in the CSDns (Figure 7B) prevents CPP. Additionally, knocking down 5-HT1B neurons globally in all serotonergic neurons has the same effect (Figure 7A). The inhibitory nature of the 5-HT1BRs implies that the reduction of 5-HT release is required to facilitate CPP. On the other hand, overexpressing 5-HT1BRs on CSDNs ensures less 5-HT release and because 5-HT release from the CSDns is required for CPP (Figure 1D), we can conclude that 5-HT levels are carefully regulated during the critical period to maintain permissive levels of E/I balance. This concentration dependent, bi-directional control allows for the maintenance of an optimal 5-HT level above or below which CPP is hindered.

An alternate mode of action of the 5-HT1BRs on serotonergic cells could be the localization of the serotonin transporters (dSERT) that promote serotonin reuptake thereby reducing extracellular 5-HT levels.^67,68,70^ Further studies are required to confirm if this mechanism holds true for serotonergic modulation of serotonin neurons during the critical period.

### Neuronal mechanism of serotonergic modulation in CPP

During the olfactory critical period, chronic activation of the OSNs by an odor, leads to activation dependent structural plasticity in the cognate glomerulus. This increase in volume can be attributed to the increase in PN and LN arborizations in an odor specific manner.^19–21,27^ A specific subpopulation of GABAergic LNs, the LN1 neurons, the adenylate cyclase *rutabaga* increases cAMP levels to promote CREB dependent gene transcription. This facilitates the structural plasticity observed in both the LNs and PNs.^20^ Functionally, it leads to an increase in inhibitory output and a decrease in excitatory PN output onto the higher order olfactory centers. This mechanism suggests a way by which the primary olfactory center modulates sensory output before it can reach the second-order olfactory centers (Figure 7C). Our observations show that serotonergic modulation integrates at multiple levels of this model to modify the output from the AL. Firstly, 5-HT release from the CSDns within this circuit is required for the structural plasticity. Additionally, 5-HT levels in the extracellular space are maintained by 5-HT1BR mediated inhibition of serotonergic neurons. Similarly, serotonergic modulation on the OSNs is tightly controlled where lower levels of 5-HT2BR expression permits structural plasticity while higher levels of 5-HT2BR expression achieved 2 days post eclosion coincides with the end of the critical period. The 5-HT7Rs on the R70A09 and therefore the LN1 neurons likely activates cAMP dependent gene transcription that ultimately leads to the increase in LN and PN arborizations resulting in glomerular volume increase (Figure 7D). Future experiments are required to examine this model at a greater detail at the cellular and molecular level using pharmacology, and electrophysiology.

In conclusion, our work provides novel insight into how 5-HT modulates structural plasticity at multiple sites of primary olfactory processing during the olfactory critical period. Specifically, we show that 5-HT directly affects the stimulus specific circuit via 5-HT2BRs on the OSNs and 5-HT7Rs on LNs which indirectly modulates GABAergic inhibition throughout the AL. Finally, we show that 5-HT release from the CSDns is carefully controlled during the critical period, disruption of which hinders CPP. This supports the view that neuromodulators affect different components of sensory processing to facilitate structural plasticity during the critical period.

## Acknowledgements

We thank Dr. Eric Horstick for helpful feedback on the manuscript and Jonathan Schenk, Clarissa Ng and Marryn Bennett for technical guidance. We also thank Dr. Ricardo Areneda for hosting parts of these experiments. This work was supported by National Institutes of Health R21DC018945-01 to Q.G. and R01 DC016293 to A.M.D. and Q.G. We thank A.E. Beaven and the UMD Imaging core facility for confocal imaging. Purchase of the Zeiss LSM 980 Airyscan 2 was supported by Award Number 1S10OD025223-01A1 from the National Institute of Health.

## Author Contributions

A.M, H.T., A.M.D and Q.G conceived the study. A.M., A.M.D. and Q.G. wrote the manuscript. A.M. J.E. and H.T. carried out most of the experiments. H.T. generated the tagged 5-HT receptor constructs and flies. A.M.D and Q.G. secured funding and supervised the project. All of the authors read and approved the manuscript.

## References

1. Mallick, A., Dacks, A.M., and Gaudry, Q. (2024). Olfactory Critical Periods: How Odor Exposure Shapes the Developing Brain in Mice and Flies. Biology 2024, Vol. 13, Page 94 13, 94. 10.3390/BIOLOGY13020094.

2. Reha, R.K., Dias, B.G., Nelson, C.A., Kaufer, D., Werker, J.F., Kolbh, B., Levine, J.D., and Hensch, T.K. (2020). Critical period regulation acrossmultiple timescales. Proc Natl Acad Sci U S A 117, 23242–23251. 10.1073/pnas.1820836117.

3. Hensch, T.K. (2005). Critical period plasticity in local cortical circuits. Nat Rev Neurosci 6, 877–888. 10.1038/nrn1787.

4. Wang, Y., Gu, Q., and Cynader, M.S. (1997). Blockade of serotonin-2C receptors by mesulergine reduces ocular dominance plasticity in kitten visual cortex. Exp Brain Res 114, 321–328. 10.1007/PL00005640/METRICS.

5. Gu, Q., and Singer, W. (1995). Involvement of Serotonin in Developmental Plasticity of Kitten Visual Cortex. European Journal of Neuroscience 7, 1146–1153. 10.1111/j.1460-9568.1995.tb01104.x.

6. Jitsuki, S., Takemoto, K., Kawasaki, T., Tada, H., Takahashi, A., Becamel, C., Sano, A., Yuzaki, M., Zukin, R.S., Ziff, E.B., et al. (2011). Serotonin Mediates Cross-Modal Reorganization of Cortical Circuits. Neuron 69, 780–792. 10.1016/J.NEURON.2011.01.016.

7. Dyck, R.H., and Cynader, M.S. (1993). Autoradiographic localization of serotonin receptor subtypes in cat visual cortex: transient regional, laminar, and columnar distributions during postnatal development. Journal of Neuroscience 13, 4316–4338. 10.1523/JNEUROSCI.13-10-04316.1993.

8. Teissier, A., Soiza-Reilly, M., and Gaspar, P. (2017). Refining the role of 5-HT in postnatal development of brain circuits. Front Cell Neurosci 11, 1–9. 10.3389/fncel.2017.00139.

9. Suri, D., Teixeira, C.M., Cagliostro, M.K.C., Mahadevia, D., and Ansorge, M.S. (2014). Monoamine-Sensitive Developmental Periods Impacting Adult Emotional and Cognitive Behaviors. Neuropsychopharmacology 2015 40:1 40, 88–112. 10.1038/npp.2014.231.

10. Higa, G.S.V., Francis-Oliveira, J., Carlos-Lima, E., Tamais, A.M., Borges, F. da S., Kihara, A.H., Shieh, I.C., Ulrich, H., Chiavegatto, S., and De Pasquale, R. (2022). 5-HT-dependent synaptic plasticity of the prefrontal cortex in postnatal development. Scientific Reports 2022 12:1 12, 1–23. 10.1038/s41598-022-23767-9.

11. Ogelman, R., Gomez Wulschner, L.E., Hoelscher, V.M., Hwang, I.-W., Chang, V.N., and Oh, W.C. (2024). Serotonin modulates excitatory synapse maturation in the developing prefrontal cortex. Nature Communications 2024 15:1 15, 1–15. 10.1038/s41467-024-45734-w.

12. Hildebrand, J.G., and Shepherd, G.M. (1997). Mechanisms of olfactory discrimination: converging evidence for common principles across phyla. Annu Rev Neurosci 20, 595– 631. 10.1146/ANNUREV.NEURO.20.1.595.

13. Hallem, E.A., and Carlson, J.R. (2006). Coding of Odors by a Receptor Repertoire. Cell 125, 143–160. 10.1016/j.cell.2006.01.050.

14. Hallem, E.A., Ho, M.G., and Carlson, J.R. (2004). The molecular basis of odor coding in the Drosophila antenna. Cell 117, 965–979. 10.1016/j.cell.2004.05.012.

15. Benton, R., Sachse, S., Michnick, S.W., and Vosshall, L.B. (2006). Atypical membrane topology and heteromeric function of Drosophila odorant receptors in vivo. PLoS Biol 4, 240–257. 10.1371/JOURNAL.PBIO.0040020.

16. Benton, R., Vannice, K.S., Gomez-Diaz, C., and Vosshall, L.B. (2009). Variant Ionotropic Glutamate Receptors as Chemosensory Receptors in Drosophila. Cell 136, 149–162. 10.1016/j.cell.2008.12.001.

17. Wilson, R.I. (2013). Early Olfactory Processing in Drosophila□: Mechanisms and Principles. Annu Rev Neurosci 36, 217–241. 10.1146/annurev-neuro-062111-150533.

18. Fabian, B., and Sachse, S. (2023). Experience-dependent plasticity in the olfactory system of Drosophila melanogaster and other insects. Front Cell Neurosci 17. 10.3389/fncel.2023.1130091.

19. Fabian, B., Grabe, V., Beutel, R.G., Hansson, B.S., and Sachse, S. (2023). Experience-dependent plasticity of a highly specific olfactory circuit in Drosophila melanogaster. bioRxiv, 2023.07.26.550642. 10.1101/2023.07.26.550642.

20. Das, S., Sadanandappa, M.K., Dervan, A., Larkin, A., Lee, J.A., Sudhakaran, I.P., Priya, R., Heidari, R., Holohan, E.E., Pimentel, A., et al. (2011). Plasticity of local GABAergic interneurons drives olfactory habituation. Proc Natl Acad Sci U S A 108, 2–10. 10.1073/pnas.1106411108.

21. Chodankar, A., Sadanandappa, M.K., Raghavan, K.V., and Ramaswami, M. (2020). Glomerulus-Selective regulation of a critical period for interneuron plasticity in the drosophila antennal lobe. Journal of Neuroscience 40, 5549–5560. 10.1523/JNEUROSCI.2192-19.2020.

22. Golovin, R.M., Vest, J., and Broadie, K. (2021). Neuron-specific FMRP roles in experience-dependent remodeling of olfactory brain innervation during an early-life critical period. Journal of Neuroscience 41, 1218–1241. 10.1523/JNEUROSCI.2167-20.2020.

23. Golovin, R.M., Vest, J., Vita, D.J., and Broadie, K. (2019). Activity-dependent remodeling of Drosophila olfactory sensory neuron brain innervation during an early-life critical period. Journal of Neuroscience 39, 2995–3012. 10.1523/JNEUROSCI.2223-18.2019.

24. Kidd, S., and Lieber, T. (2016). Mechanism of Notch Pathway Activation and Its Role in the Regulation of Olfactory Plasticity in Drosophila melanogaster. PLoS One 11, 1–26. 10.1371/journal.pone.0151279.

25. Kidd, S., Struhl, G., and Lieber, T. (2015). Notch Is Required in Adult Drosophila Sensory Neurons for Morphological and Functional Plasticity of the Olfactory Circuit. PLoS Genet 11, 1–26. 10.1371/journal.pgen.1005244.

26. Lieber, T., Kidd, S., and Struhl, G. (2011). DSL-Notch Signaling in the Drosophila Brain in Response to Olfactory Stimulation. Neuron 69, 468–481. 10.1016/j.neuron.2010.12.015.

27. Sachse, S., Rueckert, E., Keller, A., Okada, R., Tanaka, N.K., Ito, K., and Vosshall, L.B.B. (2007). Activity-Dependent Plasticity in an Olfactory Circuit. Neuron 56, 838–850. 10.1016/j.neuron.2007.10.035.

28. Hensch, T.K., and Quinlan, E.M. (2018). Critical periods in amblyopia. Vis Neurosci 35, E014. 10.1017/S0952523817000219.

29. Takesian, A.E., Bogart, L.J., Lichtman, J.W., and Hensch, T.K. (2018). Inhibitory circuit gating of auditory critical-period plasticity. Nature Neuroscience 2018 21:2 21, 218–227. 10.1038/s41593-017-0064-2.

30. Toyoizumi, T., Miyamoto, H., Yazaki-Sugiyama, Y., Atapour, N., Hensch, T.K., and Miller, K.D. (2013). A Theory of the Transition to Critical Period Plasticity: Inhibition Selectively Suppresses Spontaneous Activity. Neuron 80, 51–63.

31. Berardi, N., Pizzorusso, T., Ratto, G.M., and Maffei, L. (2003). Molecular basis of plasticity in the visual cortex. Trends Neurosci 26, 369–378. 10.1016/S0166-2236(03)00168-1.

32. Kirkwood, A. (2000). Serotonergic control of developmental plasticity. Proc Natl Acad Sci U S A 97, 1951–1952. 10.1073/pnas.070044697.

33. Kojic, L., Dyck, R.H., Gu, Q., Douglas, R.M., Matsubara, J., and Cynader, M.S. (2000). Columnar distribution of serotonin-dependent plasticity within kitten striate cortex. Proc Natl Acad Sci U S A 97, 1841–1844. 10.1073/pnas.97.4.1841.

34. Maya Vetencourt, J.F., Tiraboschi, E., Spolidoro, M., Castrén, E., and Maffei, L. (2011). Serotonin triggers a transient epigenetic mechanism that reinstates adult visual cortex plasticity in rats. European Journal of Neuroscience 33, 49–57. 10.1111/j.1460-9568.2010.07488.x.

35. Vetencourt, J.F.M., Sale, A., Viegi, A., Baroncelli, L., De Pasquale, R., O’Leary, O.F., Castrén, E., and Maffei, L. (2008). The antidepressant fluoxetine restores plasticity in the adult visual cortex. Science (1979) 320, 385–388. 10.1126/SCIENCE.1150516/SUPPL_FILE/MAYAVETENCOURT.SOM.PDF.

36. Suh, G.S.B., Wong, A.M., Hergarden, A.C., Wang, J.W., Simon, A.F., Benzer, S., Axel, R., and Anderson, D.J. (2004). A single population of olfactory sensory neurons mediates an innate avoidance behaviour in Drosophila. Nature 431, 854–859. 10.1038/nature02980.

37. Faucher, C., Forstreuter, M., Hilker, M., and De Bruyne, M. (2006). Behavioral responses of Drosophila to biogenic levels of carbon dioxide depend on life-stage, sex and olfactory context. Journal of Experimental Biology 209, 2739–2748. 10.1242/JEB.02297.

38. Schenk, J.E., and Gaudry, Q. (2023). Nonspiking Interneurons in the Drosophila Antennal Lobe Exhibit Spatially Restricted Activity. eNeuro 10. 10.1523/ENEURO.0109-22.2022.

39. Ran, F.A., Hsu, P.D., Wright, J., Agarwala, V., Scott, D.A., and Zhang, F. (2013). Genome engineering using the CRISPR-Cas9 system. Nature Protocols 2013 8:11 8, 2281–2308. 10.1038/nprot.2013.143.

40. Gratz, S.J., Ukken, F.P., Rubinstein, C.D., Thiede, G., Donohue, L.K., Cummings, A.M., and Oconnor-Giles, K.M. (2014). Highly specific and efficient CRISPR/Cas9-catalyzed homology-directed repair in Drosophila. Genetics 196, 961–971. 10.1534/GENETICS.113.160713/-/DC1.

41. Suzuki, Y., Schenk, J.E., Tan, H., and Gaudry, Q. (2020). A Population of Interneurons Signals Changes in the Basal Concentration of Serotonin and Mediates Gain Control in the Drosophila Antennal Lobe. Current Biology 30, 1110–1118.e4. 10.1016/j.cub.2020.01.018.

42. Ostrovsky, A., Cachero, S., and Jefferis, G. (2013). Clonal Analysis of Olfaction in Drosophila: Immunochemistry and Imaging of Fly Brains. Cold Spring Harb Protoc 4. doi:10.1101/pdb.prot071720.

43. Legland, D., Arganda-Carreras, I., and Andrey, P. (2016). MorphoLibJ: Integrated library and plugins for mathematical morphology with ImageJ. Bioinformatics 32, 3532–3534. 10.1093/BIOINFORMATICS/BTW413.

44. Fitzpatrick, M. (2014). Measuring cell fluorescence using ImageJ. Github. https://github.com/mfitzp/theolb/blob/master/imaging/measuring-cell-fluorescence-using-imagej.rst.

45. Gnerer, J.P., Venken, K.J.T., and Dierick, H.A. (2015). Gene-specific cell labeling using MiMIC transposons. Nucleic Acids Res 43, e56. 10.1093/NAR/GKV113.

46. Cheung, U.S., Shayan, A.J., Boulianne, G.L., and Atwood, H.L. (1999). Drosophila larval neuromuscular junction’s responses to reduction of cAMP in the nervous system. J Neurobiol 40, 1–13. 10.1002/(SICI)1097-4695(199907)40:1%3C1::AID-NEU1%3E3.0.CO;2-1.

47. Dacks, A.M., Christensen, T.A., and Hildebrand, J.G. (2006). Phylogeny of a serotonin-immunoreactive neuron in the primary olfactory center of the insect brain. Journal of Comparative Neurology 498, 727–746. 10.1002/CNE.21076.

48. Coates, K.E., Majot, A.T., Zhang, X., Michael, C.T., Spitzer, S.L., Gaudry, Q., and Dacks, A.M. (2017). Identified Serotonergic Modulatory Neurons Have Heterogeneous Synaptic Connectivity within the Olfactory System of Drosophila. Journal of Neuroscience 37, 7318–7331. 10.1523/JNEUROSCI.0192-17.2017.

49. Coates, K.E., Calle-Schuler, S.A., Helmick, L.M., Knotts, V.L., Martik, B.N., Salman, F., Warner, L.T., Valla, S. V., Bock, D.D., and Dacks, A.M. (2020). The Wiring Logic of an Identified Serotonergic Neuron That Spans Sensory Networks. Journal of Neuroscience 40, 6309–6327. 10.1523/JNEUROSCI.0552-20.2020.

50. Zhang, X., and Gaudry, Q. (2016). Functional integration of a serotonergic neuron in the drosophila antennal lobe. Elife 5, 1–24. 10.7554/eLife.16836.

51. Sun, X.J., Tolbert, L.P., and Hildebrand, J.G. (1993). Ramification pattern and ultrastructural characteristics of the serotonin-immunoreactive neuron in the antennal lobe of the moth Manduca sexta: A laser scanning confocal and electron microscopic study. Journal of Comparative Neurology 338, 5–16. 10.1002/CNE.903380103.

52. Jenett, A., Rubin, G.M., Ngo, T.T.B., Shepherd, D., Murphy, C., Dionne, H., Pfeiffer, B.D., Cavallaro, A., Hall, D., Jeter, J., et al. (2012). A GAL4-Driver Line Resource for Drosophila Neurobiology. Cell Rep 2, 991–1001. 10.1016/J.CELREP.2012.09.011.

53. Witz, P., Amlaiky, N., Plassat, J.L., Maroteaux, L., Borrelli, E., and Hen, R. (1990). Cloning and characterization of a Drosophila serotonin receptor that activates adenylate cyclase. Proceedings of the National Academy of Sciences 87, 8940–8944. 10.1073/PNAS.87.22.8940.

54. Saudou, F., Boschert, U., Amlaiky, N., Plassat, J.-L., and Hen, R. (1992). A family of Drosophila serotonin receptors with distinct intracellular signalling properties and expression patterns. EMBO J 11, 7–17. 10.1002/J.1460-2075.1992.TB05021.X.

55. Colas, J.F., Launay, J.M., Kellermann, O., Rosay, P., and Maroteaux, L. (1995). Drosophila 5-HT2 serotonin receptor: coexpression with fushi-tarazu during segmentation. Proceedings of the National Academy of Sciences 92, 5441–5445. 10.1073/PNAS.92.12.5441.

56. Qian, Y., Cao, Y., Deng, B., Yang, G., Li, J., Xu, R., Zhang, D., Huang, J., and Rao, Y. (2017). Sleep homeostasis regulated by 5HT2b receptor in a small subset of neurons in the dorsal fan-shaped body of drosophila. Elife 6. 10.7554/eLife.26519.

57. Sizemore, T.R., and Dacks, A.M. (2016). Serotonergic Modulation Differentially Targets Distinct Network Elements within the Antennal Lobe of Drosophila melanogaster. Scientific Reports 2016 6:1 6, 1–14. 10.1038/srep37119.

58. Vicario, M., Cieri, D., Vallese, F., Catoni, C., Barazzuol, L., Berto, P., Grinzato, A., Barbieri, L., Brini, M., and Calì, T. (2019). A split-GFP tool reveals differences in the sub-mitochondrial distribution of wt and mutant alpha-synuclein. Cell Death & Disease 2019 10:11 10, 1–16. 10.1038/s41419-019-2092-1.

59. Pedelacq, J.D., and Cabantous, S. (2019). Development and Applications of Superfolder and Split Fluorescent Protein Detection Systems in Biology. International Journal of Molecular Sciences 2019, Vol. 20, Page 3479 20, 3479. 10.3390/IJMS20143479.

60. Kamiyama, R., Banzai, K., Liu, P., Marar, A., Tamura, R., Jiang, F., Fitch, M.A., Xie, J., and Kamiyama, D. (2021). Cell-type-specific, multicolor labeling of endogenous proteins with split fluorescent protein tags in Drosophila. Proc Natl Acad Sci U S A 118, e2024690118. 10.1073/PNAS.2024690118/SUPPL_FILE/PNAS.2024690118.SM02.AVI.

61. Cabantous, S., Terwilliger, T.C., and Waldo, G.S. (2005). Protein tagging and detection with engineered self-assembling fragments of green fluorescent protein. Nature Biotechnology 2004 23:1 23, 102–107. 10.1038/nbt1044.

62. Romei, M.G., and Boxer, S.G. (2019). Split Green Fluorescent Proteins: Scope, Limitations, and Outlook. https://doi.org/10.1146/annurev-biophys-051013-022846 48, 19–44. 10.1146/ANNUREV-BIOPHYS-051013-022846.

63. Witz, P., Amlaiky, N., Plassat, J.L., Maroteaux, L., Borrelli, E., and Hen, R. (1990). Cloning and characterization of a Drosophila serotonin receptor that activates adenylate cyclase. Proc Natl Acad Sci U S A 87, 8940–8944. 10.1073/PNAS.87.22.8940.

64. Sizemore, T.R., Hurley, L.M., and Dacks, A.M. (2020). Serotonergic modulation across sensory modalities. J Neurophysiol 123, 2406–2425. 10.1152/JN.00034.2020/ASSET/IMAGES/LARGE/Z9K0062054960001.JPEG.

65. Root, C.M., Masuyama, K., Green, D.S., Enell, L.E., Nässel, D.R., Lee, C.H., and Wang, J.W. (2008). A Presynaptic Gain Control Mechanism Fine-Tunes Olfactory Behavior. Neuron 59, 311–321. 10.1016/J.NEURON.2008.07.003.

66. Kwon, J.Y., Dahanukar, A., Weiss, L.A., and Carlson, J.R. (2007). The molecular basis of CO2 reception in Drosophila. Proc Natl Acad Sci U S A 104, 3574–3578. 10.1073/PNAS.0700079104/ASSET/2D2BDC64-92FC-4013-B7AB-F6E2E4466D73/ASSETS/GRAPHIC/ZPQ0070753290004.JPEG.

67. Tiger, M., Varnäs, K., Okubo, Y., and Lundberg, J. (2018). The 5-HT 1B receptor - a potential target for antidepressant treatment. Psychopharmacology (Berl) 235, 1317– 1334. 10.1007/S00213-018-4872-1/TABLES/1.

68. Middlemiss, D.N., and Hutson, P.H. (1990). The 5-HT1B Receptors. Ann N Y Acad Sci 600, 132–147. 10.1111/J.1749-6632.1990.TB16878.X.

69. Brazell, M.P., Marsden, C.A., Nisbet, A.P., and Routledge, C. (1985). The 5-HT1 receptor agonist RU-24969 decreases 5-hydroxytryptamine (5-HT) release and metabolism in the rat frontal cortex in vitro and in vivo. Br J Pharmacol 86, 209–216. 10.1111/J.1476-5381.1985.TB09451.X.

70. Barnes, N.M., and Sharp, T. (1999). A review of central 5-HT receptors and their function. Neuropharmacology 38, 1083–1152. 10.1016/S0028-3908(99)00010-6.

71. Hagan, C.E., Mcdevitt, R.A., Liu, Y., Furay, A.R., and Neumaier, J.F. (2012). 5-HT1B autoreceptor regulation of serotonin transporter activity in synaptosomes. Synapse 66, 1024–1034. 10.1002/SYN.21608.

72. Larsen, B., Cui, Z., Adebimpe, A., Pines, A., Alexander-Bloch, A., Bertolero, M., Calkins, M.E., Gur, R.E., Gur, R.C., Mahadevan, A.S., et al. (2022). A developmental reduction of the excitation:inhibition ratio in association cortex during adolescence. Sci Adv 8, 8750. 10.1126/SCIADV.ABJ8750/SUPPL_FILE/SCIADV.ABJ8750_SM.PDF.

73. Levelt, C.N., and 11ubener, M. (2012). Critical-period plasticity in the visual cortex. Annu Rev Neurosci 35, 309–330. 10.1146/ANNUREV-NEURO-061010-113813/CITE/REFWORKS.

74. Erzurumlu, R.S., and Gaspar, P. (2012). Development and critical period plasticity of the barrel cortex. European Journal of Neuroscience 35, 1540–1553. 10.1111/J.1460-9568.2012.08075.X.

75. Olsen, S.R., and Wilson, R.I. (2008). Lateral presynaptic inhibition mediates gain control in an olfactory circuit. 452. 10.1038/nature06864.

76. Yaksi, E., and Wilson, R.I. (2010). Electrical Coupling between Olfactory Glomeruli. Neuron 67, 1034–1047. 10.1016/j.neuron.2010.08.041.

77. Nagel, K.I., Wilson, R.I., and Nagel, Katherine I;Wilson, R.I. (2011). Biophysical mechanisms underlying olfactory receptor neuron dynamics. Nat Neurosci 14, 208–218. 10.1038/nn.2725.

78. Liu, W.W., and Wilson, R.I. (2013). Glutamate is an inhibitory neurotransmitter in the Drosophila olfactory system. Proceedings of the National Academy of Sciences 110, 10294–10299. 10.1073/PNAS.1220560110.

79. Hong, E.J., and Wilson, R.I. (2015). Simultaneous encoding of odors by channels with diverse sensitivity to inhibition. Neuron 85, 573–589. 10.1016/j.neuron.2014.12.040.

80. Nagel, K.I., Hong, E.J., and Wilson, R.I. (2015). Synaptic and circuit mechanisms promoting broadband transmission of olfactory stimulus dynamics. Nat Neurosci 18, 56–65. 10.1038/nn.3895.

81. Gervasi, N., Tché, P., and Preat, T. (2010). PKA Dynamics in a Drosophila Learning Center: Coincidence Detection by Rutabaga Adenylyl Cyclase and Spatial Regulation by Dunce Phosphodiesterase. Neuron 65, 516–529. 10.1016/j.neuron.2010.01.014.

82. Stam, N.J., Roesink, C., Dijcks, F., Garritsen, A., Van Herpen, A., and Olijve, W. (1997). Human serotonin 5-HT7 receptor: cloning and pharmacological characterisation of two receptor variants. FEBS Lett 413, 489–494. 10.1016/S0014-5793(97)00964-2.

83. Hobson, R.J., Hapiak, V.M., Xiao, H., Buehrer, K.L., Komuniecki, P.R., and Komuniecki, R.W. (2006). SER-7, a Caenorhabditis elegans 5-HT7-like Receptor, Is Essential for the 5-HT Stimulation of Pharyngeal Pumping and Egg Laying. Genetics 172, 159–169. 10.1534/GENETICS.105.044495.

84. Omar, M., Zhou, Y., Planting, E., Parvataneni, R., and Hung, S. (2009). Combined active triangulation, morphology scheme for active shape retrieval. Sensor Review 29, 233–239. 10.1108/02602280910967648/FULL/HTML.

85. Shen, Y., Monsma, F.J., Metcalf, M.A., Jose, P.A., Hamblin, M.W., and Sibley, D.R. (1993). Molecular cloning and expression of a 5-hydroxytryptamine7 serotonin receptor subtype. Journal of Biological Chemistry 268, 18200–18204. 10.1016/S0021-9258(17)46830-X.

86. Ruat, M., Traiffort, E., Leurs, R., Tardivel-Lacombe, J., Diaz, J., Arrang, J.M., and Schwartz, J.C. (1993). Molecular cloning, characterization, and localization of a high-affinity serotonin receptor (5-HT7) activating cAMP formation. Proceedings of the National Academy of Sciences 90, 8547–8551. 10.1073/PNAS.90.18.8547.

87. Qi, Y. xiang, Jin, M., Ni, X. yang, Ye, G. yin, Lee, Y., and Huang, J. (2017). Characterization of three serotonin receptors from the small white butterfly, Pieris rapae. Insect Biochem Mol Biol 87, 107–116. 10.1016/J.IBMB.2017.06.011.

88. Dacks, A.M., Reale, V., Pi, Y., Zhang, W., Dacks, J.B., Nighorn, A.J., and Evans, P.D. (2013). A Characterization of the Manduca sexta Serotonin Receptors in the Context of Olfactory Neuromodulation. PLoS One 8, e69422. 10.1371/JOURNAL.PONE.0069422.

89. Qi, Y. xiang, Jin, M., Ni, X. yang, Ye, G. yin, Lee, Y., and Huang, J. (2017). Characterization of three serotonin receptors from the small white butterfly, Pieris rapae. Insect Biochem Mol Biol 87, 107–116. 10.1016/J.IBMB.2017.06.011.

90. Schlenstedt, J., Balfanz, S., Baumann, A., and Blenau, W. (2006). Am5-HT7: molecular and pharmacological characterization of the first serotonin receptor of the honeybee (Apis mellifera). J Neurochem 98, 1985–1998. 10.1111/J.1471-4159.2006.04012.X.

91. Pietrantonio, P. V., Jagge, C., and McDowell, C. (2001). Cloning and expression analysis of a 5HT7-like serotonin receptor cDNA from mosquito Aedes aegypti female excretory and respiratory systems. Insect Mol Biol 10, 357–369. 10.1046/J.0962-1075.2001.00274.X.

92. Röser, C., Jordan, N., Balfanz, S., Baumann, A., Walz, B., Baumann, O., and Blenau, W. (2012). Molecular and Pharmacological Characterization of Serotonin 5-HT2α and 5-HT7 Receptors in the Salivary Glands of the Blowfly Calliphora vicina. PLoS One 7, e49459. 10.1371/JOURNAL.PONE.0049459.

93. Lee, D.W., and Pietrantonio, P. V. (2003). In vitro expression and pharmacology of the 5-HT7-like receptor present in the mosquito Aedes aegypti tracheolar cells and hindgut-associated nerves. Insect Mol Biol 12, 561–569. 10.1046/J.1365-2583.2003.00441.X.

94. Vleugels, R., Lenaerts, C., Vanden Broeck, J., and Verlinden, H. (2014). Signalling properties and pharmacology of a 5-HT7-type serotonin receptor from Tribolium castaneum. Insect Mol Biol 23, 230–243. 10.1111/IMB.12076.

95. Hensch, T.K., and Fagiolini, M. (2005). Excitatory–inhibitory balance and critical period plasticity in developing visual cortex. Prog Brain Res 147, 115–124. 10.1016/S0079-6123(04)47009-5.

96. Hunter, I., Coulson, B., Pettini, T., Davies, J.J., Parkin, J., Landgraf, M., and Baines, R.A. (2024). Balance of activity during a critical period tunes a developing network. Elife 13. 10.7554/ELIFE.91599.

97. Carlos-Lima, E., Higa, G.S.V., Viana, F.J.C., Tamais, A.M., Cruvinel, E., Borges, F. da S., Francis-Oliveira, J., Ulrich, H., and De Pasquale, R. (2023). Serotonergic Modulation of the Excitation/Inhibition Balance in the Visual Cortex. International Journal of Molecular Sciences 2024, Vol. 25, Page 519 25, 519. 10.3390/IJMS25010519.

98. Kaneko, T., Macara, A.M., Li, R., Hu, Y., Iwasaki, K., Dunnings, Z., Firestone, E., Horvatic, S., Guntur, A., Shafer, O.T., et al. (2017). Serotonergic Modulation Enables Pathway-Specific Plasticity in a Developing Sensory Circuit in Drosophila. Neuron 95, 623–638.e4. 10.1016/j.neuron.2017.06.034.

