## Supplementary material for "Serotonin acts through multiple cellular targets during an olfactory critical period": Figure Supplement

1. Department of Biology, University of Maryland, College Park, MD 20742, USA.
2. Departments of Biology and Neuroscience, West Virginia University, Morgantown, WV 26505, USA.
3. Senior Author: These authors contributed equally.

**Supplementary Information**

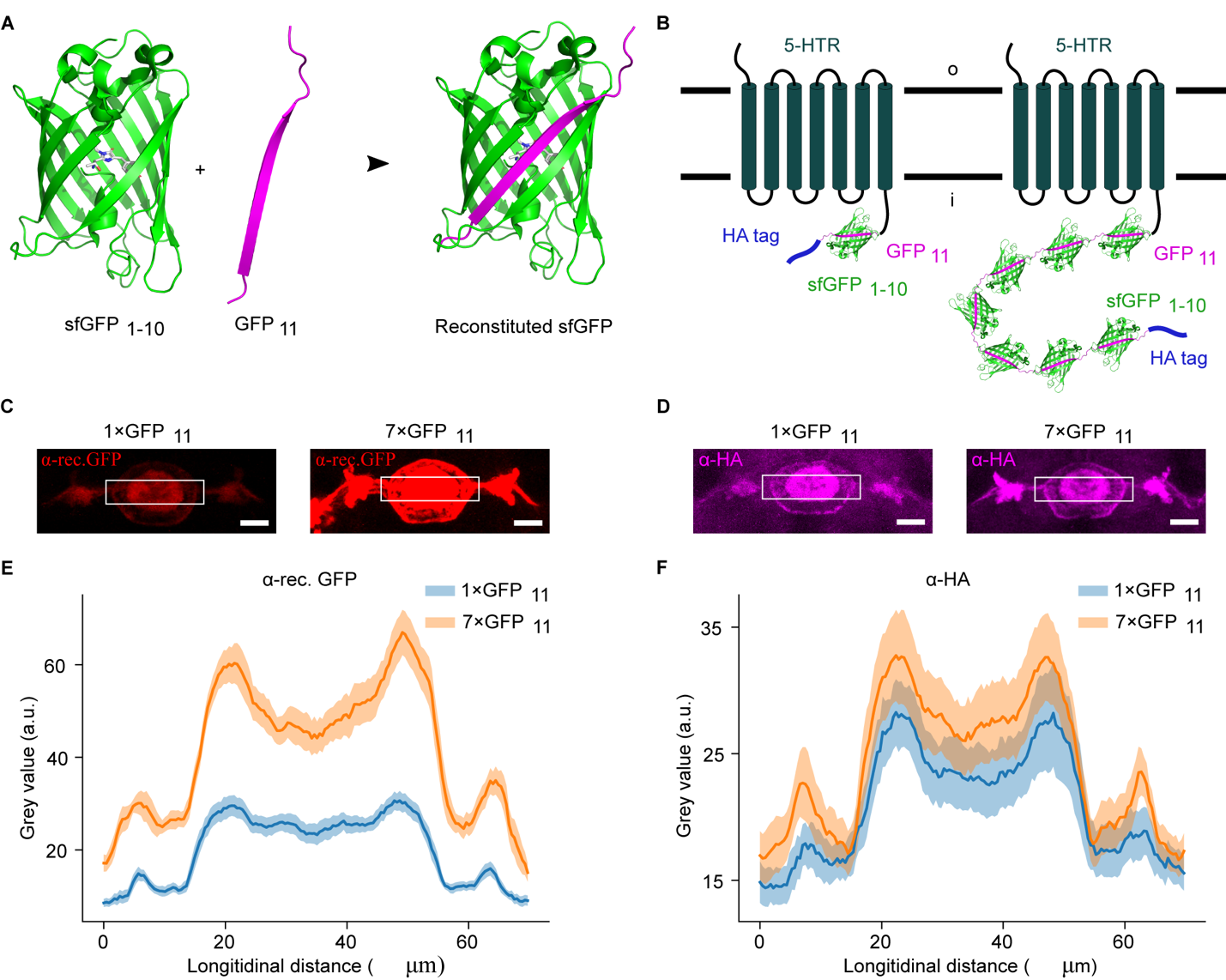

**Fig. S1. The principle of using the split-sfGFP approach to label 5-HTRs in a cell type-specific manner.**

(A) sfGFP_1-10_ (green) and GFP_11_ (magenta) reconstitute and fluoresce. PDB code: 2B3P.

(B) One or seven tandem copies of GFP_11_ fragments with an HA tag are tagged to the intracellular C terminus of 5-HTRs. sfGFP_1-10_ is introduced via the binary expression system to achieve cell type specificity.

(C) Reconstitution comparison between 5-HT7R-GFP_11_-HA (5-HT7 receptor with one copy of GFP_11_ fragment plus an HA tag fused to the C-terminus of the receptor) or 5-HT7R-7×GFP_11_-HA (5-HT7 receptor with seven copies of GFP_11_ fragments plus an HA tag fused to the C-terminus of the receptor) in the area of the ellipsoid body. sfGFP_1-10_ was brought in via 5-HT7-Gal4 (Gifted by Charles D. Nichols). The reconstitution was detected with an antibody that specifically targets the reconstituted GFP, denoted as α-rec. GFP. An 8 by 70 μm rectangle was drawn over the ellipsoid body, in which the longitudinal accumulated fluorescence intensity was collected for comparison. Scale bar, 20 μm.

(D)Similar to (C), except that an antibody that targeted the HA-tag (α-HA) was used in the immunostaining for comparison. Flies with the same genotypes as in (C) were used. Scale bar, 20 μm.

(E-F) The longitudinal accumulated fluorescence intensities collected from the rectangular regions shown in (C) and (D) were respectively plotted to compare for the α-rec. GFP channel (E) and the α-HA channel (F). In (E), N = 10 for both 1×GFP_11_ and 7×GFP_11_; in (F), N = 9 and 10 for 1×GFP_11_ and 7×GFP_11_, respectively.

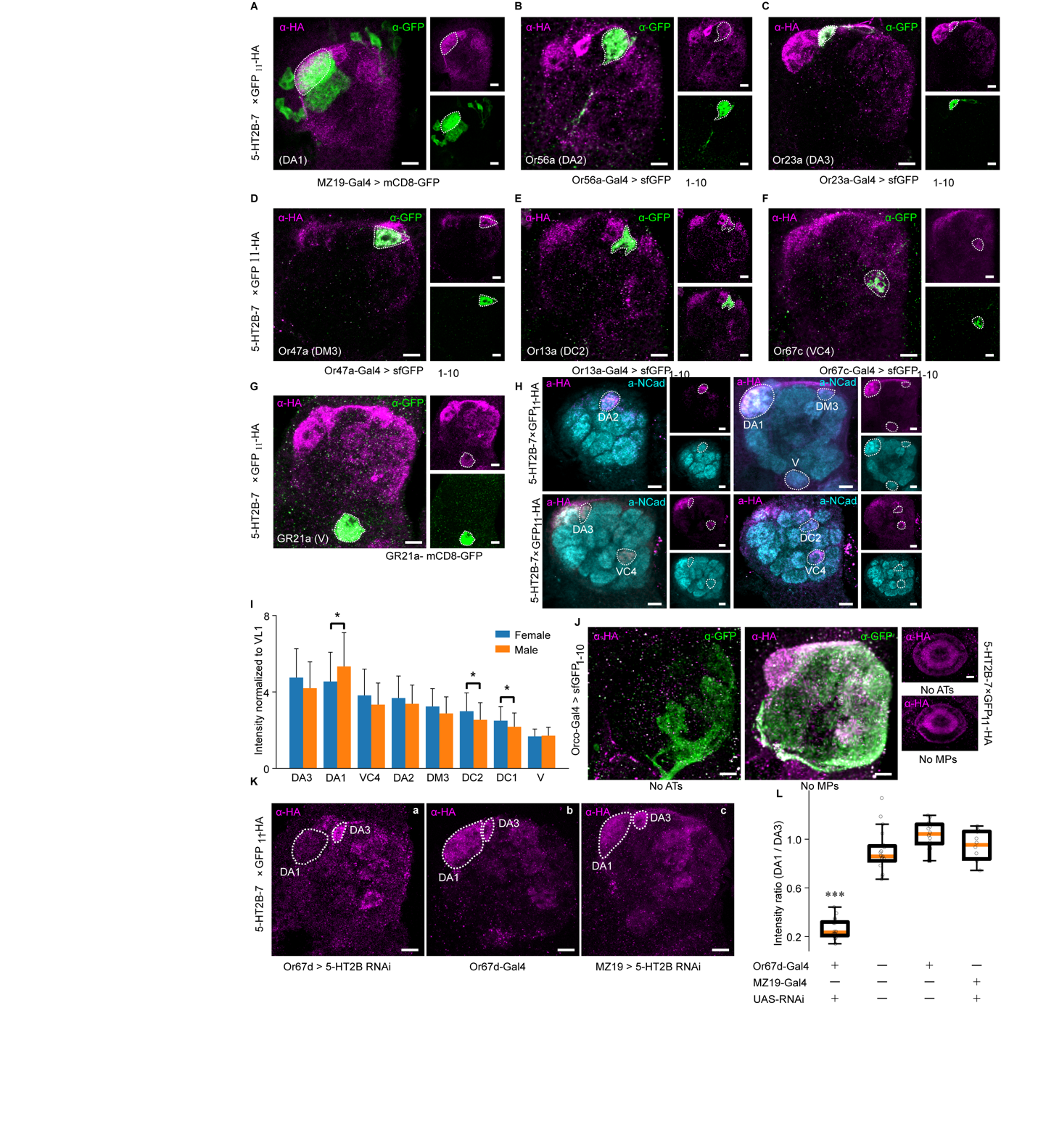

**Fig. S2. 5-HT2B receptor is expressed in multiple glomeruli in antennal lobes of Drosophila.**

(A-G) Confirmation of the identity of OSNs expressing 5-HT2B. Overlap of GFP11-HA (magenta) and sfGFP1-10 were used to identify specific glomeruli. Dotted lines demarcate the glomerulus of interest in each image. All images are organized with lateral towards left and dorsal upwards.

(H) The seven glomeruli that have a relatively high expression level of 5-HT2B revealed by the α-HA antibody (magenta) in the background staining with α-nCad (N-Cadherin, cyan). Dotted lines indicate the glomeruli of interest.

(I) In male and female brains, mean α-HA fluorescence intensity in each of the eight glomeruli was normalized to that mean signal intensity in VL1 glomerulus. Student’s t-test was performed to compare the intensities in each glomerulus between males and females. Asterisks indicate significant difference for DA1, DC1 and DC2 glomeruli, for which, p = 0.029, 0.041 and 0.027, respectively. No significant difference was found in other glomeruli. N = 43 brains for females and 44 brains for males.

(J) Four days old of female flies were severed antennae (ATs) or maxillary palps (MPs). Six days later, their brains were dissected and immunostained with α-HA (magenta) and α-GFP (green) antibodies. In addition to antennal lobes, the fluorescent state in the ellipsoid body was also examined (right).

(K) RNAi targeting 5-HT2BRs was introduced to Or67d-Gal4 labeled OSNs or MZ19-Gal4 labeled. In (a), compared to (b), the transgene for RNAi was omitted. Dotted lines indicate the DA1 glomerulus. Magenta indicates the α-HA.

(L) Under different conditions in terms of 5-HT2BR RNAi expression, the fluorescence intensity within the DA1 glomerulus in the α-HA channel was collected and normalized to that within the DA3 glomerulus. n = 9 for Or67d with RNAi, 19 for RNAi alone, 14 for Or67d-Gal4 alone and for MZ19 with RNAi.

Scale bar, 10 μm.

T**able S1. List of flies used in the supplementary figures.**

| **Figure** | **Genotype** | **Source** |
| --- | --- | --- |
| Fig. S1C – F | w1118; 10×UAS-sfGFP_1-10_/+; 5-HT7-7×GFP_11_-HA/5-HT7-Gal4 and w1118; 10×UAS-sfGFP_1-10_/+; 5-HT7-GFP_11_-HA/5-HT7-Gal4 | 5-HT7-Gal4^1^  Gift from Dr. Charles D. Nichols, LSU Health Sciences Centre in New Orleans |
| Fig. S2A | w1118 ; MZ19-Gal4, UAS-mCD8-GFP/+; 5-HT2B-7×GFP_11_-HA/+ | MZ19-Gal4 (BDSC # 34497) |
| Fig. S2B | w1118; 10×UAS-sfGFP_1-10_/+; 5-HT2B-7×GFP_11_-HA/Or56a-Gal4 | Or56a-Gal4 (BDSC #23896) |
| Fig. S2C | w1118; 10×UAS-sfGFP_1-10_/Or23a-Gal4; 5-HT2B-7×GFP_11_-HA/+ | Or23a-Gal4 (BDSC #9956) |
| Fig. S2D | w1118; 10×UAS-sfGFP_1-10_/+; 5-HT2B-7×GFP_11_-HA/Or47a-Gal4 | Or47a-Gal4 (BDSC #9982) |
| Fig. S2E | w1118; 10×UAS-sfGFP_1-10_/Or13a-Gal4; 5-HT2B-7×GFP_11_-HA/+ | Or13a-Gal4 (BDSC #9945) |
| Fig. S2F | Or67c-Gal4/W1118; 10×UAS-sfGFP_1-10_/+; 5-HT2B-7×GFP_11_-HA/+ | Or67c-Gal4 (BDSC #24856) |
| Fig. S2G | w1118; Gr21a-Mmus\Cd8a.GFP /+;5-HT2B-7×GFP_11_-HA/+ |  |
| Fig. S2H | 5-HT2B-7xGFP_11_-HA |  |
| Fig. S2J | w1118; 10×UAS-sfGFP_1-10_/Orco-Gal4; 5-HT2B-7×GFP_11_-HA/TM2 |  |
| Fig. S2K, left panel | Or67d-Gal4/ w1118; UAS-5-HT2B-RNAi/Pin; 5-HT2B-7×GFP_11_-HA/+ | Or67d-Gal4 (BDSC 23906) |
| Fig. S2K, middle panel | Or67d-Gal4/ W1118; +/+; 5-HT2B-7×GFP_11_-HA/+ |  |
| Fig. S2K, right panel | w1118;10×UAS-5-HT2B-RNAi/MZ19-GAL4; 5-HT2B-7×FP_11_-HA/+ |  |
| Fig. S2L, 1 – 4 from left to right | 1: Same to Fig. S2K, left panel  2: w1118; +/+; 5-HT2B-7×GFP_11_-HA/5-HT2B-7×GFP_11_-HA  3: Same to Fig. S2K, middle panel  4: Same to Fig. S2K, right panel |  |

**Sequence of pU6b (circular) backbone.**

GTTCGACTTGCAGCCTGAAATACGGCACGAGTAGGAAAAGCCGAGTCAAATGCCGAATGCAGAGTCTCATTACAGCACAATCAACTCAAGAAAAACTCGACACTTTTTTACCATTTGCACTTAAATCCTTTTTTATTCGTTATGTATACTTTTTTTGGTCCCTAACCAAAACAAAACCAAACTCTCTTAGTCGTGCCTCTATATTTAAAACTATCAATTTATTATAGTCAATAAATCGAACTGTGTTTTCAACAAACGAACAATAGGACACTTTGATTCTAAAGGAAATTTTGAAAATCTTAAGCAGAGGGTTCTTAAGACCATTTGCCAATTCTTATAATTCTCAACTGCTCTTTCCTGATGTTGATCATTTATATAGGTATGTTTTCCTCAATACTTCGGGGTCTTCGTAGAGTCTAGAAAACATCCCATAAAACATCCCATATTCAGCCGCTAGCATGGATGTTTTCCCAGTCACGACGTTGTAAAACGACGGCCAGTCTTAAGCTCGGGCCCCAAATAATGATTTTATTTTGACTGATAGTGACCTGTTCGTTGCAACAAATTGATGAGCAATGCTTTTTTATAATGCCAACTTTGTACAAAAAAGCAGGCTCCGCGGCCGCCCCCTTCACCACTAGAGGAGAGCAACTGCATAAGGCTATGAAGAGATACGCCCTGGTTCCTGGAACAATTGCTTTTACAGATGCACATATCGAGGTGGACATCACTTACGCTGAGTACTTCGAAATGTCCGTTCGGTTGGCAGAAGCTATGAAACGATATGGGCTGAATACAAATCACAGAATCGTCGTATGCAGTGAAAACTCTCTTCAATTCTTTATGCCGGTGTTGGGCGCGTTATTTATCGGAGTTGCAGTTGCGCCCGCGAACGACATTTATAATGAACGTGAATTGCTCAACAGTATGGGCATTTCGCAGCCTACCGTAGTGTTTGTTTCCAAAAAGGGGTTGCAAAAAATTTTGAACGTGCAAAAAAAATTACCAATAATCCAGAAAATTATTATCATGGATTCTAAAACGGATTACCAGGGATTTCAGTCGATGTGAATTCGAGAAGACCTGTTTTAGAGCTAGAAATAGCAAGTTAAAATAAGGCTAGTCCGTTATCAACTTGAAAAAGTGGCACCGAGTCGGTGCGATCCATTTTTTTGCTCACCTGTGATTGCTCCTACTCAAATACAAAAACATCAAATTTTCTGTCAATAAAGCATATTTATTTATATTTATTTTACAGGAAAGAATTACTAGTGAGCTCCAGCTTTTGTTCCCTTTAGTGAGGGTTAATTGCGCGCTTGGCGTAATCATGGTCATAGCTGTTTCCTGTGTGAAATTGTTATCCGCTCACAATTCCACACAACATACGAGCCGGAAGCATAAAGTGTAAAGCCTGGGGTGCCTAATGAGTGAGCTAACTCACATTAATTGCGTTGCGCTCACTGCCCGCTTTCCAGTCGGGAAACCTGTCGTGCCAGCTGCATTAATGAATCGGCCAACGCGCGGGGAGAGGCGGTTTGCGTATTGGGCGCTCT

Notes: U6b promotor; gRNA scaffold; The two BbsI cutting sites are located at the end of the U6b promotor and the beginning of the gRNA scaffold sequence, respectively. After cutting with Bbs I and re-ligated with the re-annealed primer pair, the chunk in between the two Bbs I sites will be replaced by the coding sequence of the gRNA spacer.

**References in Table S2**

1.	Becnel, J., Johnson, O., Luo, J., Nässel, D.R., and Nichols, C.D. (2011). The Serotonin 5-HT7Dro Receptor is Expressed in the Brain of Drosophila and is Essential for Normal Courtship and Mating. PLoS One *6*, e20800. 10.1371/JOURNAL.PONE.0020800.
